# Evolutionary conservation of sequence motifs at sites of protein modification

**DOI:** 10.1101/2022.09.15.508132

**Authors:** Shuang Li, Henrik G. Dohlman

## Abstract

Gene duplications are common in biology and are likely to be an important source of functional diversification and specialization. The yeast *Saccharomyces cerevisiae* underwent a whole genome duplication event early in evolution, and a substantial number of duplicated genes have been retained. We identified more than 3,500 instances where only one of two paralogous proteins undergoes post-translational modification despite having retained the same amino acid residue in both. We also developed a web-based search algorithm (CoSMoS.c.) that scores conservation of amino acid sequences based on 1011 wild and domesticated yeast isolates and used it to compare differentially-modified pairs of paralogous proteins. We found that the most common modifications – phosphorylation, ubiquitylation and acylation but not N-glycosylation – occur in regions of high sequence conservation. Such conservation is evident even for ubiquitylation and succinylation, where there is no established ‘consensus site’ for modification. Differences in phosphorylation were not associated with predicted secondary structure or solvent accessibility, but did mirror known differences in kinase-substrate interactions. By integrating data from large scale proteomics and genomics analysis, in a system with such substantial genetic diversity, we obtained a more comprehensive understanding of the functional basis for genetic redundancies that have persisted for 100 million years.

## Introduction

It has long been appreciated that the yeast *Saccharomyces cerevisiae* has multiple paralogous gene pairs. However, their origins were only recognized after sequencing the complete genome, the first for any eukaryotic organism. That effort, completed in 1997, revealed a whole-genome duplication event dating back roughly 100 million years (1). Of the 6,604 open reading frames (ORFs) in this organism there are 550 paralogous pairs (2–4), as annotated in the SGD YeastMine database (https://yeastmine.yeastgenome.org/yeastmine/begin.do). These duplicated genes are strongly enriched for components of the ribosome complex (4, 5) as well as of the glucose sensing pathway, including proteins involved in glycolysis and gluconeogenesis (6, 7). Thus, duplicated genes appear to be especially important for processes related to glucose utilization, and are likely to have enabled the ability of this organism to ferment glucose to ethanol even under oxygen-rich conditions.

The prevalence of paralogous genes in *Saccharomyces cerevisiae* led to broader questions about the evolutionary and selective pressures that favor their retention. Some discussions of gene paralogs have focused on their potential contributions to genetic robustness and phenotypic plasticity (8). Robustness refers to a number of different mechanisms that stabilize phenotype against genetic or environmental perturbations (e.g. changes in glucose or oxygen availability). An extreme example of robustness is where one of the genes is inactivated and the remaining copy provides enough of the original function to compensate for the loss and ensure survival. In support of this model, several studies in yeast have found that about a third of duplicate gene pairs exhibit negative epistasis (9–12), meaning that deleting both copies produces a significantly larger defect than that of the individual deletions. An alternative scenario is where the activity of a duplicated gene product is temporarily disabled in response to changing environmental circumstances, for example through substrate inhibition or feedback phosphorylation. In that case the remaining paralog might compensate for the loss by modifying its activity through transcriptional reprograming (13), changes in protein stability, or redistribution within the cell (8, 14, 15). Another feature of duplicated genes is phenotypic plasticity, which refers to the potential of new genes to evoke new phenotypes, new metabolic functions, increased biological complexity and – ultimately – the emergence of new species ((16, 17); reviewed in (18–21)). These processes are not mutually exclusive; that is to say, paralogs may allow for adaptation of a given species to a broader set of environmental circumstances and at the same time help to accelerate genetic evolution.

Underlying any changes in cellular fitness are the biochemical changes that occur within the cell. Most prominently, changes in protein function are driven by chemical modifications such as phosphorylation, glycosylation, acylation and peptidylation (e.g. ubiquitylation). A subset of these changes occurs dynamically and allows the cell to react quickly to internal and external perturbations. Here we sought to determine how paralogs in yeast differ with regard to posttranslational modifications. This was done with the expectation that chemical changes confer functional differences to otherwise structurally similar proteins, and may account for the retention of duplicated gene pairs. To facilitate our analysis, we built a web-based tool that allows detailed sequence comparisons across 1012 *Saccharomyces cerevisiae* strains, and used this to investigate how sequence conservation near the sites of modification could account for the observed differences between paralogous protein pairs. While our analysis is limited to duplicated genes in yeast, the approach could be adapted to study other closely-related protein isoforms in other organisms, and thereby reveal some of the evolutionary forces responsible for their existence.

## Results

### I. Development of CoSMoS.c

Our initial objective was to determine if closely related proteins undergo distinct chemical modifications, and if those changes are associated with unique amino acid sequence motifs near the sites of modification. To that end we analyzed sites of post-translational modifications in S288C and compared the sequence of this strain with 1011 additional isolates of *Saccharomyces cerevisiae* (22). These represent multiple clades from multiple geographical regions and from multiple sources in the wild, in clinical settings, in the laboratory, or used commercially in the dairy and brewing industries. In comparison to many other genomes of interest, including humans, there is much greater genetic diversity within the species *Saccharomyces cerevisiae*. In contrast to other yeasts, including commonly studied species such as *Candida albicans* and *Schizosaccharomyces pombe*, *Saccharomyces cerevisiae* has a substantial number of paralogous gene pairs.

To begin our analysis we first performed Multi-Sequence Alignment (using Clustal Omega 1.2.4) for 5,776 open reading frames shared among the 1012 strains (see Materials and Methods) (22). We then used these alignments to identify and analyze specific sites of interest (described below), which were the foundation for subsequent conservation score calculations.

We then created an interactive website, CoSMoS.c. (Conserved Sequence Motifs in Saccharomyces cerevisiae, https://shiny-server-dept-yeast-cosmos.apps.cloudapps.unc.edu/) that allows users to identify, either by sequence motif or by position within the sequence, and score the conservation of aligned regions of any protein (Figures 1, S1 and S2). To address multiple potential applications we included five widely-used algorithms to calculate conservation scores (see Materials and Methods for details): Shannon Entropy, which reports the average level of uncertainty (or “information” or “surprise”) inherent in the possible outcomes of the variable and thereby quantifies amino acid diversity at a given position (23); Stereochemically Sensitive Entropy, which is based on Shannon Entropy but groups amino acids into nine categories based on similarities in their physiochemical properties (24); PhyloZOOM, which weights evolutionary relatedness on top of chemical identity (22, 25); Jensen-Shannon Divergency (JSD), which compares amino acid frequencies with the background distribution, assuming no evolutionary constraint, and thereby emphasizes selection pressure rather than chemical similarity (26); and Karlin Substitution Matrix, which quantifies the likeliness of observed substitutions, rather than the chemical or biological properties of a given amino acid (27). Thus, each algorithm considers a different aspect of amino acid sequence, which when used together provides a more comprehensive representation of protein conservation.

**Figure 1.**
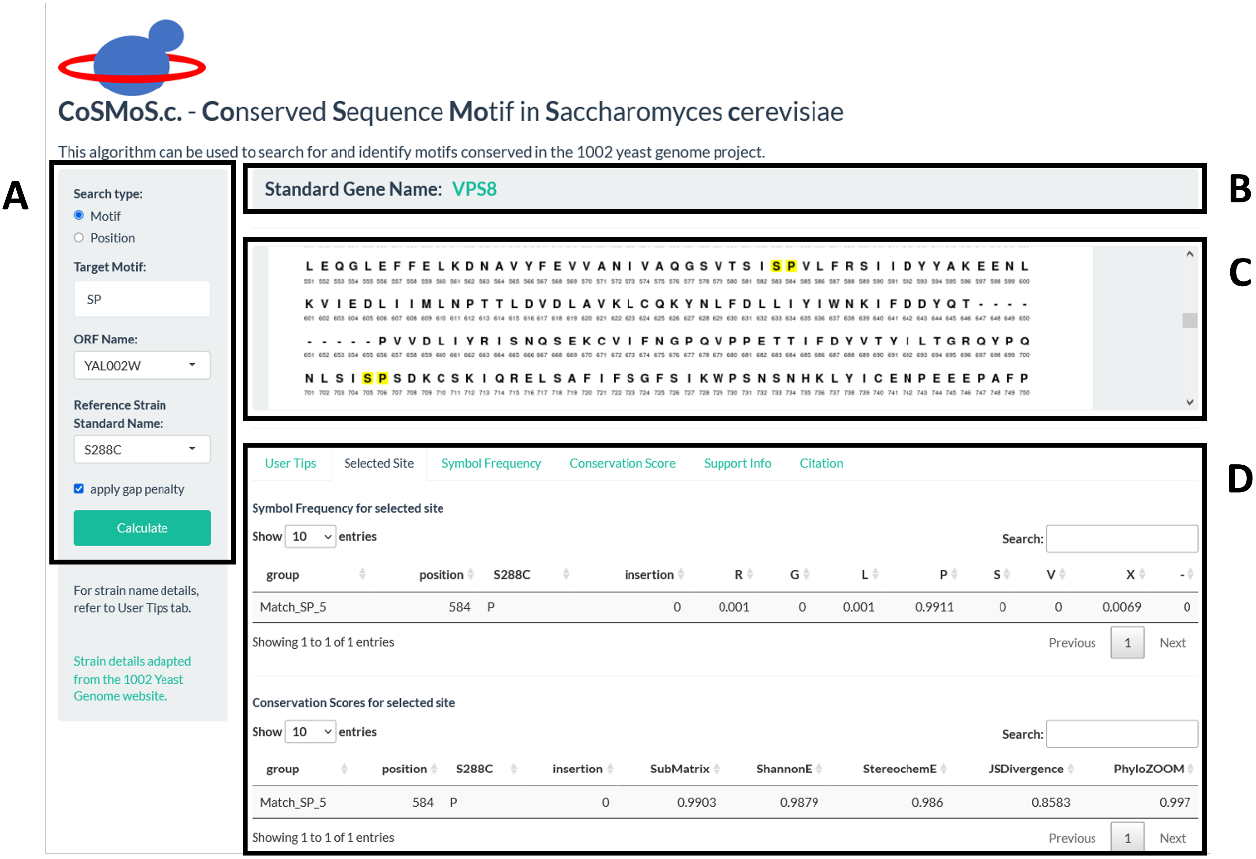
The landing page for the CoSMoS.c. website contains input and output display panels. (A) The input panel contains options for search type, ORF name, reference strain name, and whether or not to apply a gap penalty. The reference strain is used for the PhyloZOOM algorithm. The gap penalty, when applied, decreases the score if there are non-standard amino acids at the target site. The search type can be either by motif or by position within the sequence. The output panel contains three sections (top, middle, bottom). (B) The top section shows the standard gene name and contains a hyperlink to the corresponding page of the SGD website. (C) The middle section shows the numbered amino acid sequence. A position or motif that matches the input is indicated by highlighting. The highlighted region is interactive; that is to say, clicking on a highlighted amino acid will display the Frequency Table and Conservation Score table under the “Selected Site” tab (D). The bottom panel contains five additional tabs. “User Tips” provides brief instructions on how to input data and what to expect in terms of output. “Support Info” provides details on how to interpret the output statistics and a hyperlink to a detailed user manual. “Citation” tab shows how to reference the CoSMoS.c. website. “Symbol Frequency” and “Conservation Score” provide statistics for all matched sites, as detailed in Figures S1 and S2, respectively.

### II. Statistical analysis using CoSMoS.c

Proteins can undergo any of dozens of post-translational modifications, and these changes in chemical structure can have important biological consequences. Soon after the completion of the yeast genome sequencing project, many large-scale investigations of protein modifications were conducted using mass spectrometry. These experiments have shown that the vast majority of proteins are modified. Other proteins are likely to be expressed and/or modified, but may have been missed because of low abundance or because of difficulties with protein isolation or peptide detection.

One potential, and particularly powerful, use of CoSMoS.c. is to determine whether specific modifications occur in regions of high sequence conservation. To that end we aligned the sequences of paralogous protein pairs and selected those amino acid sites that are the same in both paralogs, but where only one of the two sites is modified. We then used CoSMoS.c. to calculate the conservation scores for those sites of interest and their flanking regions. Our underlying hypothesis was that if there are sequences that favor a given modification, such sequences are likely to flank the site of interest, and those sequences are likely to be conserved.

To maximize statistical power we focused on the five most common modifications in *Saccharomyces cerevisiae:* phosphorylation, with 38,684 occurrences present in 4,120 ORFs (62.39%), ubiquitylation, with 5,299 occurrences in 1,872 ORFs (28.35%), monoacetylation, with 968 occurrences in 333 ORFs (5.04%), N-glycosylation, with 587 occurrences in 239 ORFs (3.62%), and succinylation, with 577 occurrences in 356 ORFs (5.39%) (Table 1 and Dataset S1), as annotated for strain S288C in the SGD database (https://yeastmine.yeastgenome.org/yeastmine/begin.do). All of these modifications can affect protein activity, location or protein-protein interactions.

**Table 1.** Most-common modifications in the complete proteome (all gene counts) of *Saccharomyces cerevisiae* were obtained from annotated lists in the SGD database and assigned to 550 paralogs (paralog gene counts). Instances where a modified residue is not conserved (FALSE) or conserved (TRUE), regardless of modification status, in its paralog were identified after aligning paralogous protein sequences using the Needleman algorithm (needle conserved).

|  |  | all gene counts | paralog gene counts |  |  |  |
| --- | --- | --- | --- | --- | --- | --- |
|  |  |  | needle conserved |  |  |  |
| Modification Type | amino acid | Total | FALSE | TRUE | Total | conserved |
| phosphorylation | S | 30029 | 4997 | 3290 | 8287 | 39.70% |
| phosphorylation | T | 7723 | 1329 | 797 | 2126 | 37.49% |
| phosphorylation | Y | 932 | 147 | 109 | 256 | 42.58% |
| phosphorylation | S/T/Y | 38684 | 6473 | 4196 | 10669 | 39.33% |
| ubiquitylation | K | 5299 | 632 | 1160 | 1792 | 64.73% |
| (mono)acetylation | K | 968 | 49 | 306 | 355 | 86.20% |
| N-glycosylation | N | 587 | 48 | 91 | 139 | 65.47% |
| succinylation | K | 577 | 77 | 471 | 548 | 85.95% |

Phosphorylation is the addition of a phosphate group from ATP to the hydroxyl group of serine, threonine or tyrosine. Protein phosphorylation was first documented by Krebs and Fischer, who showed that this modification is responsible for the interconversion of active (“a”) and inactive (“b”) forms of the enzyme glycogen phosphorylase (reviewed in (28)). Their investigations led to the establishment of the first hormonal cascade of successive enzymatic reactions, kinases acting on kinases, initiated by cAMP and leading to glycogenolysis (28). In *Saccharomyces cerevisiae*, phosphorylation of serine, threonine and tyrosine has been documented 30,029, 7,723, and 932 times, respectively (Table 1). The abundance of phosphorylated tyrosines was unexpected given that in yeast there are no dedicated tyrosine kinases and only a small number of dual-specificity kinases. We did not include rare non-canonical events, including phosphorylation of other amino acids or events annotated as “dephosphorylation” or “autophosphorylation”.

First discovered in the late 1970s, ubiquitylation entails the conjugation of a 76 amino acid protein, ubiquitin, to lysine residues in substrate proteins. This process is mediated by three distinct enzymes (E1, E2 and E3), the last of which defines substrate specificity and the timing of the modification (29). The first use of mass spectrometry to map a protein ubiquitylation site was done for the yeast G protein Gpa1 (30). In this example, a protein complex containing the E3 Cdc53 and F-box adapter Cdc4 is necessary and sufficient for poly-ubiquitylation of Gpa1, resulting in degradation by the proteasome protease complex (31, 32). Another E3, Rsp5, is necessary and sufficient for mono-ubiquitylation (33). The monoubiquitylated protein is then internalized by a cascade of ubiquitin-binding domain proteins and degraded in the vacuole (31, 34).

The two acylation reactions, representing the addition of a succinyl or acetyl group to lysine. The first documented, and best characterized, substrates are histones. In this instance, Lys acetylation disrupts DNA binding, owing to the resulting charge reversal from +1 to −1, and alterations in chromatin structure (35). Acetylation is carried out by lysine acetyltransferases and reversed by histone deacetylases. Past computational studies suggest that these enzymes prefer to modify substrates at sites rich in Lys, Ser, Thr, Gly, and Ala (36).

Finally, N-glycosylation is the attachment of an oligosaccharide moiety to the amide nitrogen of asparagine. This modification usually occurs on proteins destined for the cell surface, either as secreted or as integral membrane proteins. In contrast to the other modifications, N-glycosylation is considered irreversible and occurs during, rather than after, protein synthesis. Oligo-saccharyltransferases transfer a preassembled oligosaccharide from a lipid-linked donor to Asn residues within glycosylation acceptor “sequons”: Asn-X-Thr/Ser/Cys, where X is any residue other than Pro. Not all sequons are glycosylated however, reflecting the importance of other primary and secondary structural features (37, 38). Once conjugated, these N-linked oligosaccharides undergo further processing, the products of which are specifically recognized by ER-localized lectins. Collectively these events facilitate proper protein folding and transport to the cell surface (Reviewed in (37)).

We next identified pairs of sites in which one of the two proteins is known to be modified and the amino acid residue is the same in both (“site of interest”). These sites were identified by aligning sequences using the Needleman algorithm (39). We did not consider substitutions of any kind even if they are potentially modified (e.g. serine for threonine). Then we sought to determine whether the modifications occurred within a region of high amino acid conservation, or conserved sequence motif. To that end we examined the flanking region of each site of interest using three methods of analysis and five different scoring algorithms within CoSMoS.c (see Materials and Methods). We are not inferring any causal relationship between a particular modification and the sequence context of that modification, particularly since different substrates may undergo the same chemical reaction but are carried out by different modifying enzymes. Further experimentation is needed to determine if a particular sequence helps to direct a specific enzyme to the site of modification, or if a given modification favors retention of sequences that are functionally compatible with a given post-translational modification.

The first method, which we call Symmetric Average Score, considers the conservation scores for sets of amino acids upstream and downstream of the site of interest. That is to say, we obtained mean scores for sets of amino acids that include, and bracket on both sides, the sites of interest. These scores are represented as mean1 (3 amino acids), mean2 (5 amino acids), mean3 (7 amino acids) and mean4 (9 amino acids) (Figure S3A). We then compared the Symmetric Average Score for each paralog pair. To determine whether the modified target site had a higher context sequence conservation than that of the unmodified paralog we first separated the scores into five groups, one for each modification type. We then performed two statistical tests: the Distribution Mean Test, which determines whether target protein conservation score distribution is significantly larger than that of its paralog, and the Paralog Pairing Test, which tests whether the pairing structure confers an advantage for the target proteins (Figure 2). We also list how these two tests are expected to perform under all possible scenarios in regard to the relationships between modified target proteins and unmodified paralogs (see Materials and Methods).

**Figure 2.**
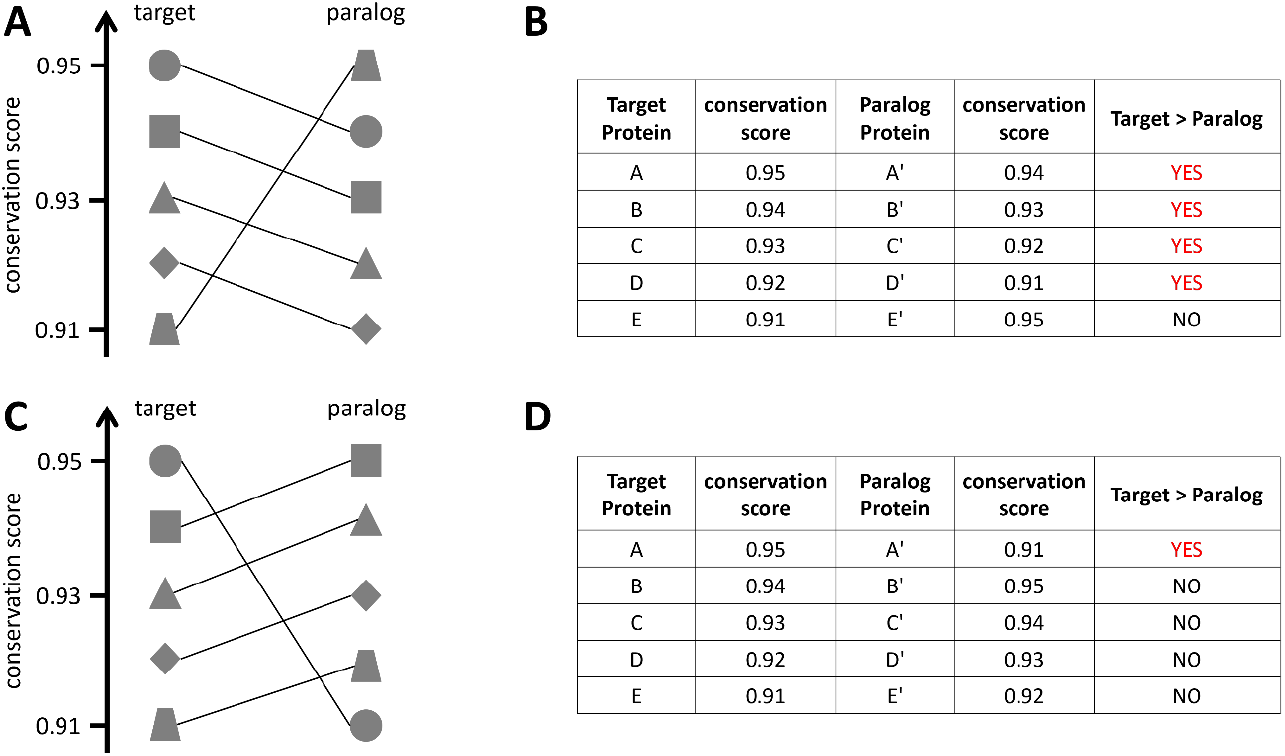
Schematic illustrating how pairing structure can advantage or disadvantage target proteins. In all panels, target and paralog proteins have the same conservation scores, but the pairing structure is such that target proteins have higher scores in four out of five instances (A and B) or target proteins have higher scores in only one out of five instances (C and D).

The second method we call One-sided Average Score. In contrast to Symmetric Average Score, which simultaneously considers sequences on both sides, this alternative method considers up to four amino acids either upstream or downstream (but not both) of the site of interest (Figure S3B). As with the first method we compared the score for each paralog pair, separated the scores into the five modification types, and performed the same statistical tests on the score distribution.

The third method, which we call Chemical Similarity Average Score, calculates the mean conservation score based on chemical classifications assigned to residues immediately adjacent to a site of interest. That is to say, we obtained a mean score for amino acids, comprised of the site of interest and the residue immediately before (“meanb1”) or immediately after (“meana1”). We then placed each of these amino acids into five separate bins based on the chemical classification of the adjoining residue, as follows: aliphatic (G,A,V,L,I,M,P), aromatic (F,Y,W), polar uncharged (S,T,C,N,Q), acidic (E,D) and basic (K,R,H). We then compared the modified target proteins and their unmodified paralogs, by calculating the average of meanb1 and of meana1, for each classification. Because the residues of a given target and its paralog could fall into two different chemical classifications, the paralog pairing structure cannot be maintained in this analysis. Therefore, we only applied the Mann-Whitney-Wilcoxon Test to compare the means for the target and paralog distribution of a certain chemical classification. In this way we determined whether there was a significant difference in conservation scores between modified target proteins and unmodified paralogs for each chemical group. Thus, we could determine if the occurrence of a specific modification depends on the physiochemical properties of nearby amino acids.

### III. Analysis of sequence conservation near sites of post-translational modification

Having established our analytical approach, we next sought to apply it to specific post-translational modifications. To that end we focused on five major modifications, representing those with more than 500 documented occurrences in the proteome of *Saccharomyces cerevisiae*, as annotated in the SGD YeastMine database (https://yeastmine.yeastgenome.org/yeastmine/begin.do): phosphorylation, N-glycosylation, monoacetylation, succinylation and ubiquitylation. More specifically, we sought to identify patterns of sequence conservation near each modification site, and to determine how any such sequence motifs differ depending on the type of modification.

We first applied the Mann-Whitney-Wilcoxon (Distribution Mean) and Monte Carlo Simulation (Paralog Pairing) tests to the Symmetric Average Score. Figures 3 and 4 show the adjusted p values (Benjamini-Hochberg, same method used for all adjusted p values) for each of the five modifications, one per column, as applied to all four sequence lengths, one per row. For the Distribution Mean Test we found significant differences (adjusted p < 0.05) in conservation scores comparing modified target proteins and unmodified paralog proteins, for all modifications except N-glycosylation (Figure 3 and Dataset S2). We also found significant differences for all sequence lengths, from one to four amino acids, flanking the site of interest. This suggests that there is a functional relationship between these modifications and their adjoining sequences. Once again, we are not inferring any mechanistic relationship between any modification and the sequence context of that modification.

**Figure 3.**
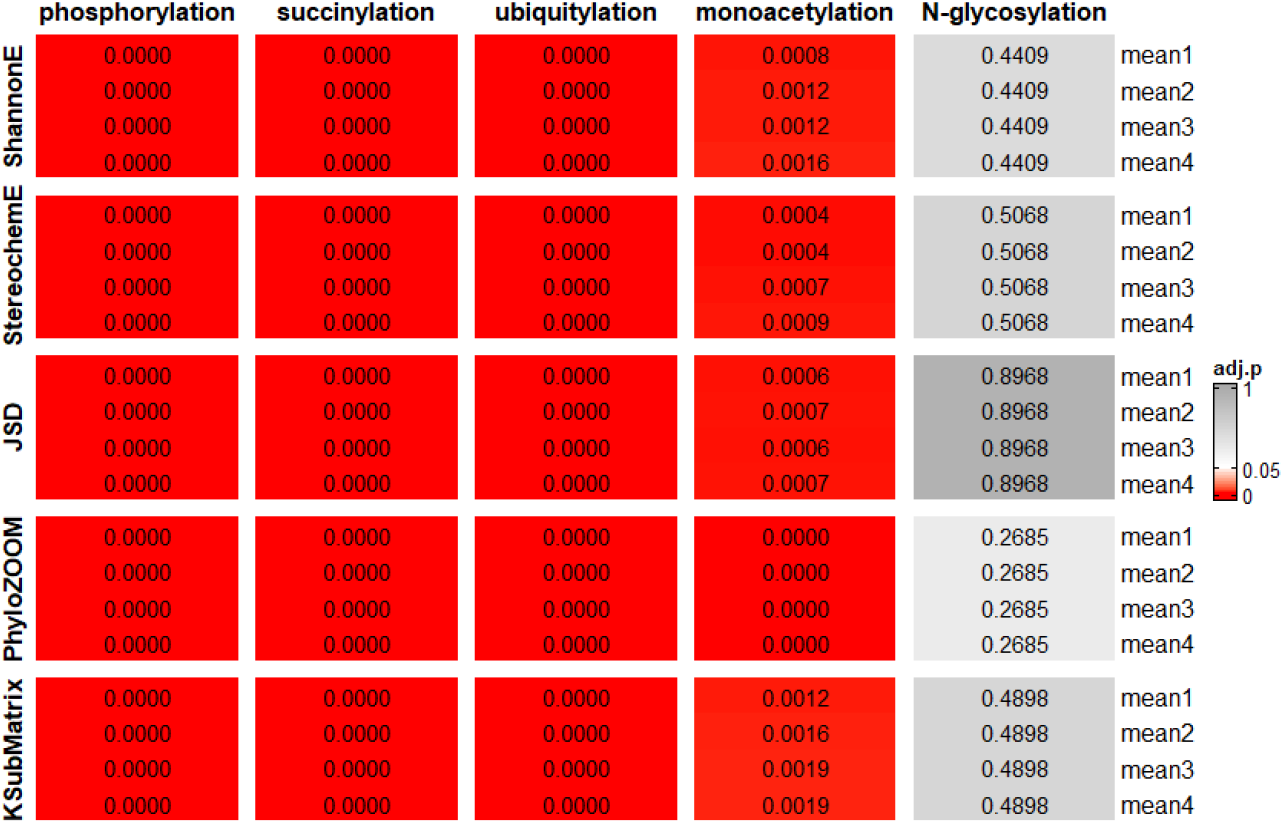
Results of Distribution Mean Test for Symmetric Average Score. Displayed is a heatmap of adjusted p values for all five algorithms with different flanking range (mean1 to mean4, rows) for each modification type (columns). Gray, p > 0.05; white, p = 0.05; red, p < 0.05.

We then applied the Paralog Pairing Test to each modification type using all five algorithms (Figure 4 and Dataset S3). Once again, conservation scores for phosphorylation and ubiquitylation were significantly different for all four sequence lengths and for all five algorithms. Succinylation had borderline adjusted p values for mean1 scores obtained using Shannon Entropy and Karlin Substitution Matrix.

**Figure 4.**
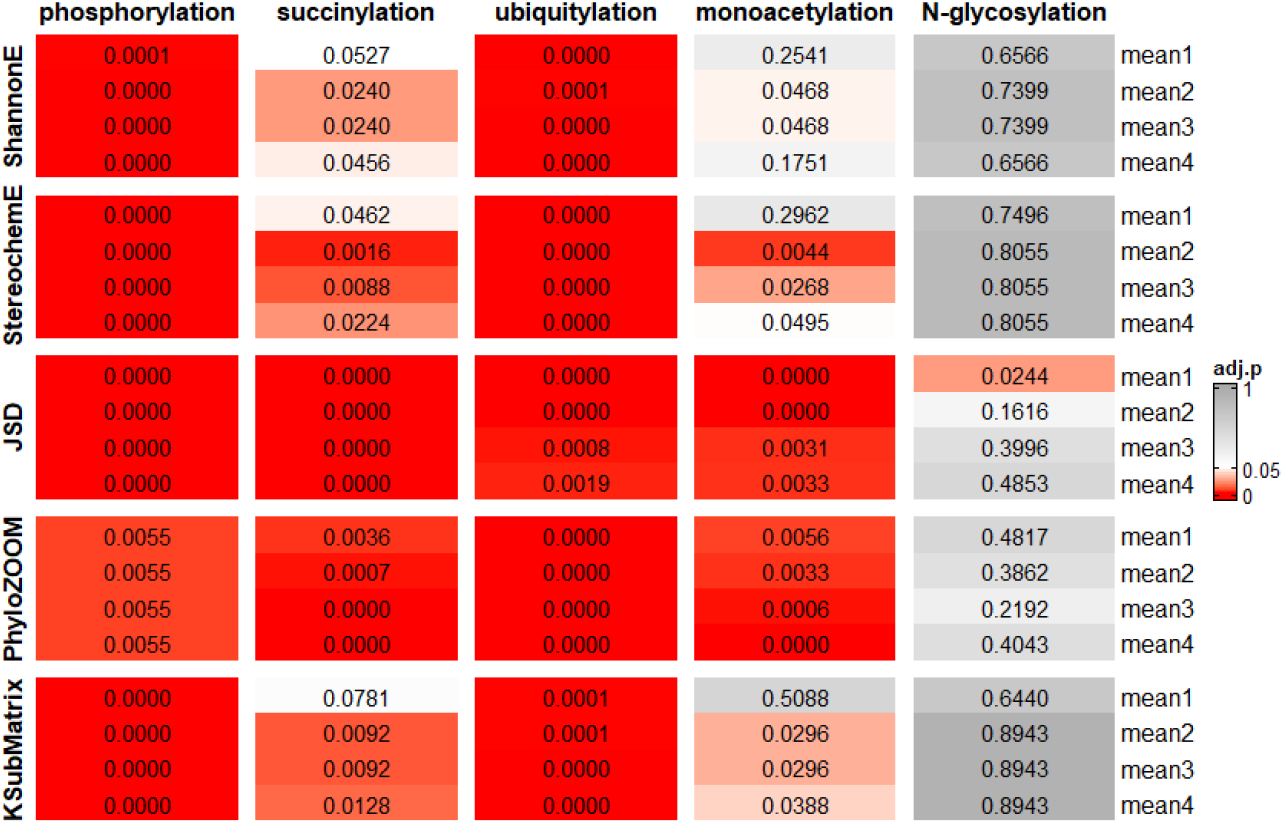
Results of Paralog Pairing Test for Symmetric Average Score. Displayed is a heatmap of adjusted p values for all five algorithms with different flanking range (mean1 to mean4, rows) for each modification type (columns). Gray, p > 0.05; white, p = 0.05; red, p < 0.05.

Monoacetylation had high adjusted p values for three of five mean1 scores (all but JSD and PhyloZOOM). JSD reports how much we expect the amino acid sequence to change assuming no evolutionary constraint. The JSD value is greater for modified targets than unmodified paralogs. Therefore, the significant adjusted p value suggests that selection pressure is greater near modified sites than unmodified sites. In contrast to the other algorithms, PhyloZOOM penalizes mutations according to phylogenetic distance. We found that for mean1 of monoacetylation the modified target proteins are more likely to have a PhyloZOOM score that is equal to or greater than that of unmodified paralogs. Accordingly, the sequences of unmodified paralogs exhibit substitutions in strains closely related to S288C, while modified proteins harbor substitutions in distantly related strains. In summary, JSD indicates that there is selection pressure to ensure the conservation of sequences flanking sites of modification, and these forces are relaxed for corresponding sites in unmodified paralogs. PhyloZOOM indicates the selection pressure for modified regions is greater in strains most closely related to S288C. Such differences are less likely to be detected when using other algorithms such as Shannon Entropy. Finally, for most algorithms monoacetylation had a significant adjusted p value for mean2, mean3 and mean4. Based on these data we infer that the pairing structure helps to amplify differences between target proteins and their paralogs, and these differences are evident for phosphorylation, ubiquitylation, succinylation and monoacetylation, but not for N-glycosylation. We obtained similar results using the One-sided Average Score, which considers up to four amino acids either upstream or downstream of the site of interest, but not both. For both Distribution Mean Test and Paralog Pairing Test, adjusted p values were significant for segments upstream and downstream of the sites of phosphorylation, succinylation, ubiquitylation, and monoacetylation, but not N-glycosylation (Figures S4, S5 and Datasets S4, S5).

Finally, we determined the Chemical Similarity Average Score, which calculates the mean conservation score based on chemical classifications assigned to amino acid residues. We used the Mann-Whitney-Wilcoxon Test to compare modified target proteins and unmodified paralogs for each chemical classification. Once again, we observed significant differences for all modifications except N-glycosylation (Figure 5 and Dataset S6). In particular, we found high conservation of aliphatic residues flanking sites of phosphorylation and ubiquitylation, and immediately after sites of succinylation. In addition, we found high conservation of basic residues flanking sites of ubiquitylation, and of polar uncharged residues flanking sites of phosphorylation. Notably, we observed a significant difference for all five amino acid classifications upstream of the sites of phosphorylation. This result indicates the importance of conservation at this position, one that is independent of the chemical properties of the amino acids at that position. One possibility is that protein kinases share the ability to recognize amino acids immediately before the site of modification, but that dependence may differ for different kinases. More broadly, we conclude that a subset of amino acids is conserved near sites of protein phosphorylation, succinylation, and ubiquitylation. This is true even when there is no obvious ‘consensus site’ for a given modification. This pattern of conservation could indicate regions that are functionally important and happen to undergo post-translational modification. Alternatively, flanking sequences could favor recognition by the enzymes that confer these modifications, or disfavor recognition by the enzymes that remove them. Further studies are needed to establish a cause-effect relationship between each modification, modifying enzymes, and the sequence context of the various modifications.

**Figure 5.**
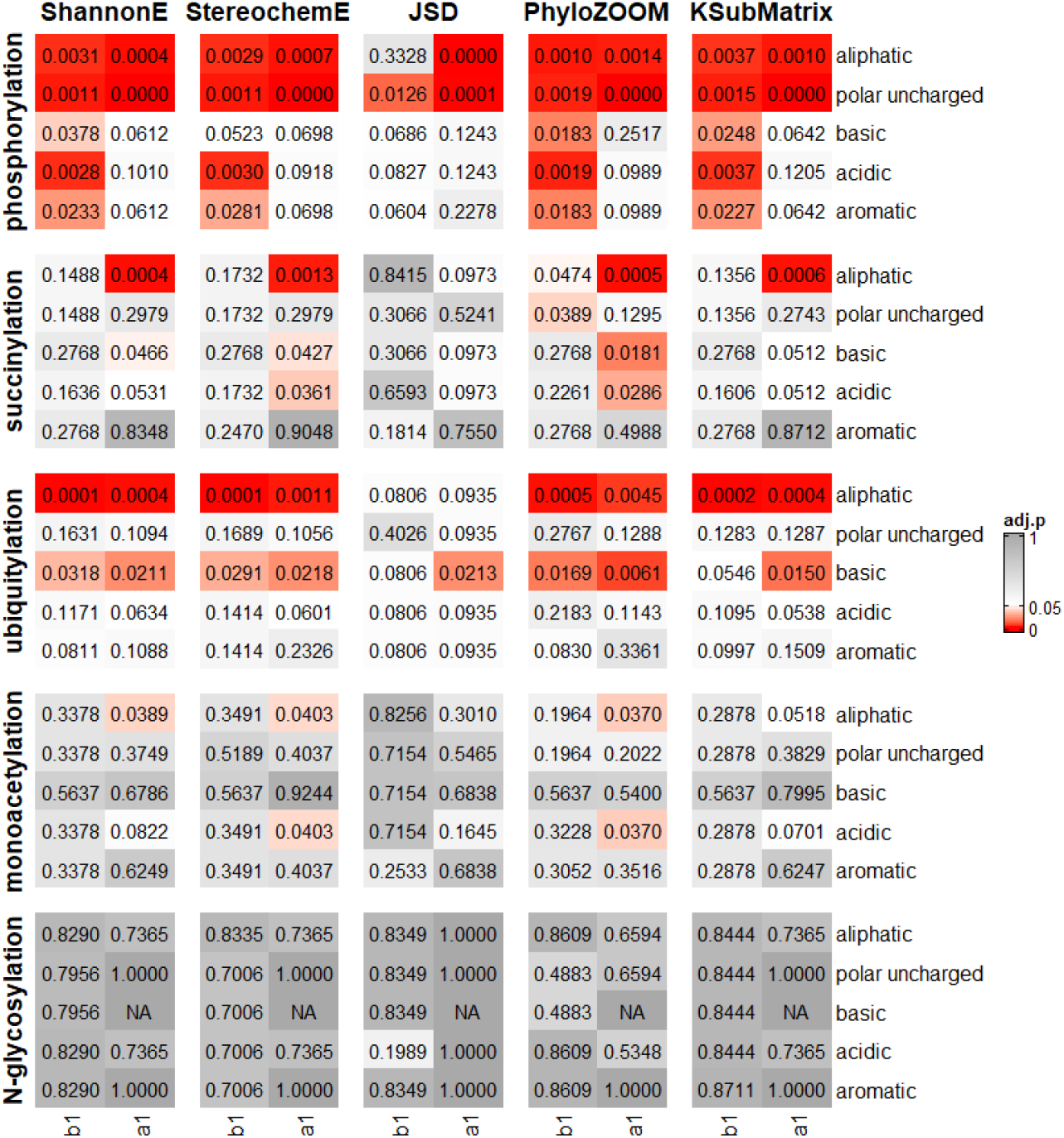
Results for Chemical Similarity Average Score. Displayed are adjusted p values for all five algorithms (column pairs) and all five modifications (rows). For each modification type, amino acid categories immediately adjacent to the site of interest (first column, upstream-b1; second column, downstream-a1) are plotted separately. Gray, p > 0.05; white, p = 0.05; red, p < 0.05.

### IV. Analysis of sequence motifs near sites of post-translational modification

We next examined specific features of amino acids flanking the sites of modification, and paid particular attention to motifs unique to modified target proteins (Table 2). In this analysis we focused on phosphorylation, since it is well established that many protein kinases recognize specific amino acid residues at positions near the site of phosphorylation (40, 41) (https://services.healthtech.dtu.dk/service.php?NetPhos-3.1; (https://scansite4.mit.edu/#home). To that end we used the Chi-square test, to determine whether the amino acid distribution for a specific site differs between modified targets and unmodified paralogs, and performed post-hoc analysis to determine which amino acid underlies this difference (see Materials and Methods) (42–44). Not surprisingly, most of the statistically significant differences were observed for serine phosphorylation, which is by far the most common. In this instance we detected significant or marginally significant enrichment of arginine at position b3, glycine at position b1 and proline at position a1. Conversely, certain amino acids were disfavored at positions b3 (asparagine) and a1 (serine, threonine, tyrosine and lysine). There is good agreement with the sequence features identified in our analysis and that of a recent analysis using combinatorial peptide library screening methods to determine substrate specificity determinants for 303 purified serine-threonine kinases in humans.^1^ Nearly half of the kinases could be assigned to sequence motifs enriched either in basic residues at the b3 and b2 positions or a proline at the a1 position.

**Table 2.** Analysis of sequence motifs near sites of post-translational modification. Shown are flanking amino acids before (b3, b1) and after (a1) favored (green) or disfavored (red) in modified target proteins, as compared to their unmodified paralog. Dark colors, adjusted p < 0.05; light colors, adjusted p < 0.10.

| modification | relative position | amino acid | relative abundance | adjusted p value |
| --- | --- | --- | --- | --- |
| phosphorylated S | b3 | R | Target > Paralog | < 0.0001 |
| phosphorylated S | b3 | N | Target < Paralog | 0.0007 |
| phosphorylated S | b1 | G | Target > Paralog | 0.0641 |
| phosphorylated S | a1 | S | Target < Paralog | 0.0442 |
| phosphorylated S | a1 | K | Target < Paralog | 0.0262 |
| phosphorylated S | a1 | T | Target < Paralog | 0.0262 |
| phosphorylated S | a1 | Y | Target < Paralog | 0.0949 |
| phosphorylated S | a1 | P | Target > Paralog | < 0.0001 |
| phosphorylated T | a1 | P | Target > Paralog | 0.0675 |

For the other types of modifications we did not observe any significant differences between target and paralog (Table 2). However, we did observe some differences in amino acid abundances, compared to that of BLOSUM62, which is denoted as the background amino acid frequencies approximating those with no selection pressure (see Materials and Methods) (45). Therefore, significant difference from BLOSUM62 distributions can be viewed as being constrained by evolution or having functional importance. For sites of monoacetylation, positively-charged amino acids (lysine and arginine) were rarely present at the preceding (b1) position (Figure S6A). For succinylation the subsequent (a1) site was enriched for lysine, aspartic acid and cysteine (Figure S6B). As with monoacetylation, the b1 position lacked arginine (Figure S6C). N-glycosylation, like phosphorylation, occurred in regions with an abundance of serines and threonines; these residues were almost exclusively found at the a2 position, in accordance with the known consensus site for N-glycosylation, as noted above (Figure S6D).

### V. Analysis of structural and interactome differences at sites of phosphorylation

Above we identified several thousand instances where one of two paralogous proteins undergoes a unique post-translational modification, and developed an algorithm that scores the conservation of sequences in modified targets and their unmodified paralogs. We then considered two possible mechanisms by which these differences might arise. First, we considered secondary structure. Modifications such as phosphorylation often occur in regions that lack obvious secondary structure; previous estimates indicate that 80% or more of phosphorylation sites lie within disordered regions of proteins, particularly proteins expressed in the cytoplasm (46, 47). The lack of structure is thought to provide increased accessibility to modifying enzymes. Second, we considered protein interaction partners. Any such differences could account for the differences in phosphorylation reported here.

In order to determine the role of secondary structure, we mapped phosphorylation sites onto predicted protein three-dimensional structures available through AlphaFold (48, 49). This algorithm provides highly accurate structure prediction by incorporating novel neural network architectures and training procedures based on the evolutionary, physical and geometric constraints of protein structures. We downloaded all available AlphaFold protein structure predictions for *Saccharomyces cerevisiae* and used STRIDE to assign residue solvent accessible area and secondary structure for each amino acid (50). We then matched secondary structure predictions to each of the modified target sites and the corresponding unmodified paralog pairs (Dataset S7). As shown in Figure S7, the most frequently observed structures for the target-paralog pairs are Coil-Coil (1363), AlphaHelix-AlphaHelix (712), Turn-Turn (400), and Strand-Strand (243), accounting for more than 80% of the total (3343). Therefore, there are no substantial differences in secondary structure type comparing targets and paralogs. We then examined each paralog pair for differences in secondary structure length or residue solvent accessible area. For these four most common secondary structure pairings, the distribution of the difference between target vs paralog of secondary structure length (Figure S7A), and residue solvent accessible area (Figure S7B) all centered near 0 with sides that were not significantly skewed in either direction. We conclude that secondary structure and residue solvent accessible area are unlikely to account for the functional differences between target and paralog.

We next considered the possibility that the paralogous proteins have distinct binding partners and these differential interactions could account for the different modifications observed. To that end, we interrogated the Yeast KID-kinase interaction database (http://www.moseslab.csb.utoronto.ca/KID/index.php) (51), representing a total of 31,155 documented interactions between protein kinases and potential substrates. Of these, 7,142 are interactions of protein kinases with paralogs. Because of limitations of the data source, we cannot assign a specific kinase to any particular amino acid modification. Therefore, we performed our analysis with all 550 paralogous protein pairs instead of individual sites in the modified target vs unmodified paralog. For each of the paralog pairs, we counted the number of kinases that interacted with both proteins (double interaction) and the number that interacted with only one of the two proteins (single interaction). We then calculated the single interaction ratio as the number of kinases with the single interaction divided by the sum of kinases with either single or double interactions. From this analysis, we identified 250 paralogous protein pairs (out of 550) having a single interaction ratio of 1 (Figure 6 and Dataset S8), meaning that these kinase interactions are exclusively with one of the two paralogous proteins, but not both. In comparison, if kinases interacted equally with both paralogous proteins, we would have a single interaction ratio of 0. These results indicate that most kinases regulate one or the other of the protein paralogs. They suggest further that differential modifications reported here may be the result of differential interactions with modifying enzymes.

**Figure 6.**
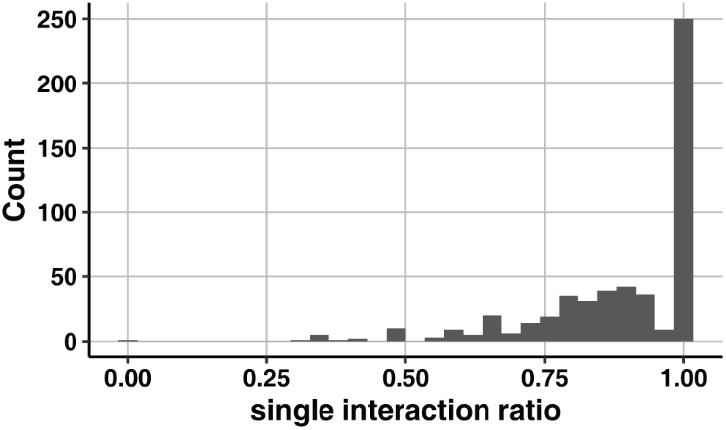
Histogram of single interaction ratio for all paralog pairs. Single interaction ratio, the number of kinases that interact with one but not both paralogs, divided by the sum of the number of kinases that interact with one or both paralogs.

## Discussion

Here we have identified several thousand instances where one of two paralogous proteins undergoes a unique post-translational modification. Source datasets were organized and analyzed using our custom algorithm (CoSMoS.c.) that quantifies sequence conservation, in an automated fashion, across 1012 unique strain isolates. By integrating multiple complementary datasets in this way, we determined that post translational modifications are associated with nearby differences in amino acid sequences, and that these modifications occur in regions of comparatively high sequence conservation. Such conservation is evident even for ubiquitylation and succinylation, where there is no established ‘consensus site’ for modification. We postulate that sequence differences confer differences in post-translational modifications, and that these modifications allow otherwise similar proteins to be differentially regulated.

Our analysis of differentially-modified pairs of paralogous proteins revealed that the most common modifications – phosphorylation, ubiquitylation and acylation but not N-glycosylation - occur within regions of high sequence conservation. In this regard, our analysis builds on past efforts to elucidate the sites and sequence determinants of protein post-translational modification. The first to be identified was the phosphorylation site in glycogen phosphorylase, in 1959 (52). In this instance, a single phosphoserine was identified after digestion with a series of proteases, followed by paper electrophoresis and end-group analysis of ^32^P-labeled peptides. That effort led to the concept of “consensus sequences” for protein modification, and was first validated using synthetic peptide substrates for cAMP dependent protein kinase (53–55). It would be another four decades before mass spectrometry would be used to survey the phosphoproteome of yeast *S. cerevisiae* (56). Subsequent studies have documented nearly 40,000 sites of phosphorylation, comprising more than 60% of the yeast proteome (Table 1). This is likely to be an underestimate, since some proteins will have been missed due to inefficient protein extraction, poor protein expression, or a low stoichiometry of phosphorylation.

Soon after the discovery of dynamic protein phosphorylation, Phillips et al. first reported the modification of histones by acetylation (57). It would be another 25 years before the first acetylation site was mapped however, in this case by epitope mapping with a monoclonal antibody specific for acetylated α-tubulin (58, 59). Acetylation-specific antibodies were later used to identify 388 acetylation sites in 195 proteins (60). Mass spectrometry was then used to identify 3,600 acetylation sites in 1,750 proteins (61). The first report of protein ubiquitylation was in the 1970s, but it was not until 2002 that a ubiquitylation site, in this case for the yeast G protein Gpa1, was mapped by mass spectrometry (30). This approach was later adopted to map more than 1000 sites of ubiquitylation in yeast (62), and is now used routinely in large scale proteomics analysis. A particular advantage of working with *Saccharomyces cerevisiae* is the existence of several hundred paralogous gene pairs, most with well-annotated functions. However, their persistence over millions of years has remained a puzzle. Duplicated genes are inherently unstable, and one or the other copy is likely to accumulate mutations and become a pseudogene or be eliminated entirely (63–65). Despite these countervailing forces, a substantial number of paralogs has been retained, presumably because these duplication events confer some fitness advantage to the organism. One potential benefit of gene duplication is to increase protein expression and metabolic flux (66, 67). In support of this dosage amplification model, many paralogous gene pairs exhibit substantial functional redundancy, as indicated by the high frequency of shared protein interaction partners (68, 69). On the other hand, it has been difficult to identify any growth or metabolic phenotypes following deletion of individual paralogs (9–12, 70). Even a combined deletion of 13 single paralogs, each involved in the conversion of glucose to ethanol, revealed no defect with respect to gene expression, the formation of glycolytic products or growth in a variety of conditions (6). Similarly, a combined deletion of 24 single paralogs associated with 40S ribosomes led to only mild loss of translation activity and cellular fitness (71). One possibility is that differences in fitness exist, but are evident only under highly specialized (non-laboratory) circumstances. Another possibility is that deletion of a given paralog leads to a compensating change in the remaining paralog, through alterations in gene transcription (13, 17), protein half-life (72) or protein abundance (73). Given this complication, and in contrast to most prior studies which examined the functional consequences of individual gene deletions, we focused our analysis on events that occur in an otherwise wildtype background.

Another potential benefit of gene duplication is the acquisition of new functions, or neofunctionalization. In this scenario, one copy might retain the original function while the second is free to acquire novel functions that are genetically favored (3). In support of the model, an analysis of published fitness data revealed that combined deletion of paralogous pairs in *Saccharomyces cerevisiae* exhibit a stronger defect than that of the corresponding singletons in *Schizosaccharomyces pombe*, which did not undergo the same whole genome duplication event (74). The expanded functionality of duplicates appears to include a larger number of protein binding partners and increased presence within multi-protein complexes (75–80). We propose that the many of the 3,500+ unique modifications – those occurring in one of two duplicated proteins – are the consequence of differential binding to protein kinases (as shown in Figure 6), ubiquitin ligases, acyl transferases, and other regulatory enzymes.

A third potential outcome of gene duplication is subfunctionalization, where the function of a single ancestral protein becomes distributed among two descendant proteins. It has been argued, based on analysis of genetic and protein interactions as well as directed evolution studies, that subfunctionalization is associated with whole-genome duplications while neofunctionalization is most characteristic of small-scale duplications (12, 75, 79). By this mechanism, duplicates can be maintained in the genome by acquiring reciprocal loss-of-function mutations, such that both duplicates become necessary to perform the combined functions of a common ancestor. These functions are likely to include distinct regulation by phosphorylation (81, 82) and changes in catalytic activity (83) or subcellular localization (84, 85). However, nearly all prior studies of subfunctionalization have focused on differences in transcriptional regulation (7, 68, 70, 86–91). In contrast, we focused our analysis on changes that occur later, through post-translational modifications of the encoded proteins.

By working with yeast we circumvent many of the challenges associated with more complex biological systems. Nevertheless, our investigations in yeast could help to explain the prevalence of seemingly redundant protein isoforms in humans. To use one example from our own prior studies (92), there are hundreds of G proteins and G protein-coupled receptors, many of which appear to be functionally interchangeable (93). For example, of the nine adrenergic receptors, all of which bind to epinephrine (adrenaline), three activate the Gi subfamily of G proteins. Gi consists of a heterotrimer comprised of one (out of three) α subunits, one (out of four) β subunits and one (out of twelve) γ subunits. Gi in turn regulates nine isoforms of adenylyl cyclase (94), all of which produce the second messenger cAMP. This single example represents 3,888 possible combinations of effector, G protein heterotrimer, and receptor. Based on our findings, we postulate that post-translational modifications may be an important source of neo- or sub-functionalization in this and other cell signaling pathways.

In conclusion, we have shown that closely related proteins can undergo very different modifications. Whereas some paralogous pairs have partitioned their functions through changes in protein sequence alone, others are likely to have acquired new roles through post-translational modifications. These observations provide a possible explanation for the retention of functionally similar proteins throughout evolutionary history.

## Materials and Methods

### Multi-Sequence Alignment

Multi-sequence alignments were performed using the reference strain S288C as well as 1011 other strains provided by the “1002 Yeast Genome” website (http://1002genomes.u-strasbg.fr/files/) (22). Gene sequences comprising 6015 non-redundant ORFs were downloaded from the allReferenceGenesWithSNPsAndIndelsInferred.tar.gz file. To simplify our analysis the 239 intron-containing genes were not considered, leaving 5776 ORFs. The S288C and “1002” datasets were combined for a total of 1012 strain sequences. These gene sequences were translated into protein sequences with the ‘translate’ function in seqinr package in R (https://cran.r-project.org/web/packages/seqinr/index.html). The translated protein sequences were used as input for Clusto Omega (version 1.2.4), with the arguments ‘--seqtype=Protein --infmt=fasta --outfmt=fasta --guidetree-out=user_defined_routes --use-kimura --iter=2 --force’, to produce multi-sequence alignments. The output was 5776 files, one for each protein, each containing an alignment for the 1012 strains.

For each site of modification the Needleman-Wunsch global alignment was performed to identify corresponding regions in each paralogous protein pair (39). The function ‘pairwiseAlignment’ in the R package ‘Biostrings (v3.14)’ and the following arguments were used: type=‘global’,substitutionMatrix = ‘BLOSUM62’, gapOpening=10,gapExtension=0.5, scoreOnly=F.

### Conservation score calculation

To calculate conservation scores we applied five commonly used algorithms. Each algorithm considers a different aspect of amino acid sequence, which when used together provides a more comprehensive representation of protein conservation. The code used to calculate the scores is available for download (https://github.com/Shuang-Plum/YeastMotifConserv).

In CoSMoS.c., ShannonEntropy was calculated as described previously (23), and was defined as (1-entropy) to be consistent with other scores. It reports the average level of uncertainty (or “information” or “surprise”) inherent in the possible outcomes of the variable, and thereby quantifies amino acid diversity at a given position.

Stereochemically Sensitive Entropy was calculated as for ShannonEntropy except that amino acids were grouped based on the rules ([‘V’,’L’, ‘I’,’M’], [‘F’.’W.’Y’], [‘S’,’T’], [‘N’,’Q’], [‘H’,’K’,’R’], [‘D’,’E’], [‘A’,’G’], [‘P’], [‘C’]), as described previously (24). Amino acids within the same group are treated as a single entity. Variation is only considered when an amino acid from one of these groups is replaced with an amino acid from another group. Therefore, Stereochemically Sensitive Entropy quantifies physiochemical similarity rather than chemical identity.

Jensen-Shannon Divergency (JSD) was calculated as first applied in (26), and as summarized as Capra07 (see Table 4 in (95)). JSD quantifies the similarity between two probability distributions. In CoSMoS.c. we used BLOSUM62 (BLOcks SUbstitution Matrix62) as the background amino acid distribution, which approximates the distribution of amino acid sites subject to no evolutionary pressure. This matrix is built using sequences with less than 62% similarity (sequences with ≥ 62% identity are clustered). BLOSUM62 is the default matrix for protein BLAST. This is also the designated background distribution in Capra07 and was shown to have broad applicability (26). Therefore JSD reports how much we expect the amino acid sequence to change assuming no evolutionary constraint. If the observed changes differ substantially from expectation (BLOSUM62), this suggests the presence of selection pressure and functional importance. Thus JSD emphasizes selection pressure rather than chemical similarity.

PhyloZOOM was calculated using the ‘zoom’ method, as described in (25). It is based on Shannon Entropy and uses a prebuilt phylogenetic tree based on the biallelic SNPs of the 1012 strains (22). This algorithm integrates strain phylogeny information into the Shannon Entropy conservation score calculation and in this way corrects for relatedness among strains. It imposes a high penalty if a mutation occurs in a comparison strain closely related to the reference strain (in our case, S288C), and imposes a low penalty if it occurs in a distantly related comparison strain. Therefore, PhyloZOOM weights evolutionary relatedness on top of chemical identity.

Karlin Substitution Matrix was calculated with Karlin Normalization as described in (27). The range of the score is (−1,1). This was then re-ranged to (0,1) to be consistent with other scores. The Karlin Substitution Matrix algorithm emphasizes the probability of amino acid substitutions. Karlin Substitution Matrix sums the weights set for each possible substitution pair based on a background distribution, which sets the rules for how a substitution is penalized. CoSMoS.c. uses BLOSUM62 to be consistent with protein BLAST, wherein a rare substitution is penalized more than a common substitution. However, the background distribution can be substituted, for example with PAM30 (96), if another set of penalty rules is preferred. Therefore, Karlin Substitution Matrix quantifies the likeliness of observed substitutions, rather than quantifying chemical or biological properties of a given amino acid.

Thus, all scores were normalized to (0,1) with 0 being random and 1 being perfectly conserved. If gap penalty was applied, Conservation Score was calculated as Score*(non-gap amino acids percentage). Gaps were defined as noncanonical amino acids X,B,Z or “–” produced from multi-sequence alignment. Gap penalty is inherent in the Karlin Substitution Matrix algorithm; therefore, no additional gap penalty was applied.

### Statistical tests for sequence conservation scores

We performed two different statistical tests because the underlying distribution has a pairing structure for each modified target protein and its unmodified paralog. One possibility is that the target protein score distribution is much larger than that of its paralog, and the distributions do not overlap (Figure S8A). In this situation the pairing structure does not matter and the target protein score is unambiguously larger than that of its paralog (Figure S8D). In this instance we applied a one-sided, paired Mann-Whitney-Wilcoxon Test (97), which determines whether the target protein conservation score distribution is significantly larger than that of unmodified paralog, without assuming that they follow a normal distribution. Hereafter we refer to this as Distribution Mean Test.

A second possibility is that modified target proteins and unmodified paralogs have conservation score distributions that overlap substantially (Figure S8B), but with a pairing structure such that the target protein score is usually higher than that of its paralog (Figures 2A, 2B and S8D). Another possibility is that the pairing structure could disadvantage the target protein, such that the target protein score is usually lower than that of its paralog (Figures 2C and 2D). To test whether the pairing structure matters we applied the Monte Carlo Simulation. We first calculated the percentage of pairings for which the modified target protein scores were greater than that of their unmodified paralogs, using established paralog gene pairs (“authentic pairs”). We then shuffled the pairings of target proteins and paralogs and calculated the percentage as before and repeated this 10,000 times. Lastly, we calculated the frequency for which “authentic pairs” was greater than that of the simulations. This method allowed us to determine if the authentic pairing structure confers an advantage for the target proteins. Hereafter we refer to this as “Paralog Pairing Test”.

A third possibility is that a modified target protein and unmodified paralog have conservation score distributions that overlap partially (Figure S8C). In this case, the Distribution Mean Test will reveal whether the mean difference is statistically significant. Under these same circumstances the results of the Paralog Pairing Test might not be significant, which would indicate that the pairing structure is not contributing to the advantage of the target protein (Figure S8D).

If the mean conservation score distribution of modified target proteins is substantially larger than that of the unmodified paralogs, such that the distributions do not overlap, the result of the Distribution Mean Test would undoubtedly be statistically significant. In this situation the result of the Paralog Pairing Test will not be significant, as no matter how the pairing is structured all target proteins will have a higher mean conservation score than that of the paralogs. Therefore, in this scenario, the target protein has a significantly higher mean conservation score than its paralog, but the pairing structure does not contribute to the significance of that difference.

If the mean conservation score distribution of the modified target proteins is larger but overlapping with that of the unmodified paralogs, the results of both the Distribution Mean Test and the Paralog Pairing Test could be statistically significant. In this situation, while the mean distribution of target proteins is significantly larger than that of the unmodified paralogs, the pairing structure could also contribute to the significance of the difference. If the mean conservation score distribution of the modified target protein is substantially overlapping with that of the unmodified paralog, the results of the Paralog Pairing Test could still be statistically significant, while the results of the Distribution Mean Test will undoubtedly not be significant. Although the distributions are similar, the pairing structure can still result in the modified target protein having a significantly larger mean conservation score than that of the unmodified paralog (See Figures 2 and S8).

### Analysis of sequence motifs near sites of post-translational modification

We made two comparisons, 1) whether there are significant differences in amino acid distribution, for a specific site, between target and paralog, and 2) whether the target or paralog distributions are significantly different from the background frequencies based on BLOSUM62. Amino acid frequencies for each of the four positions upstream and downstream of each target modification site, and the corresponding regions of the unmodified paralog protein were counted. The Chi-square test (chisq.test in stats package of R, with arguments: simulate.p.value = T, B=10000) was used to determine whether the categorized distributions, i.e. amino acid counts, were significantly different comparing 1) target proteins and paralogs, and 2) either target proteins or paralogs with BLOSUM62. To further determine which category (amino acid) contributes to the significant differences if any, post-hoc analysis was performed using the standardized residuals (stdres from chisq.test results, using pnorm to get the cumulative probability at the value of stdres, then using Benjamini-Hochberg for multi-comparison correction).

### Analysis of post-translation modifications

Modifications in the proteome of *Saccharomyces cerevisiae* were obtained from annotated lists in the SGD database (https://yeastmine.yeastgenome.org/yeastmine/begin.do) on September 27, 2021, and were assigned to 550 paralog sequences also obtained from SGD YeastMine on July 14, 2020.

### Analysis of structural motifs near sites of post-translational modification

AlphaFold protein structure prediction for *Saccharomyces cerevisiae* was downloaded from AlphaFold website (https://alphafold.ebi.ac.uk/download) (48, 49). The uncompressed.pdb files generated by AlphaFold were used as input data for protein secondary structure assignment with STRIDE as a stand-alone program (July 7th, 2022) (http://webclu.bio.wzw.tum.de/stride/) (50). STRIDE assignments of secondary structure and residue solvent accessible area were extracted from “Detailed secondary structure assignment section”. These secondary structure predictions were then matched to each of the modified target and unmodified paralog pairs. Secondary structure length is defined as the number of amino acids with an uninterrupted stretch of secondary structure, for a given site of modification or for the corresponding site of the unmodified paralog. For each modified target and unmodified paralog pair, the difference between the residue solvent accessible area and secondary structure length was calculated and presented as a boxplot for each set of target and paralog secondary structures (e.g. turn-turn, turn-coil, etc.).

### Analysis of kinase-substrate interactions

Kinase and substrate interaction data were downloaded from the Yeast KID-kinase interaction database (http://www.moseslab.csb.utoronto.ca/KID/index.php) on May 27, 2022 (51). A total of 31,155 interactions were downloaded for all substrates and including all types of experimental evidence. The dataset included 127 kinases in total. For each of the 550 paralog pairs, the number of kinases that interacted with both proteins (double interaction) and the number that interacted with only one of the two proteins (single interaction) was counted. The single interaction ratio was then defined as (number of kinases with single interaction)/(number of kinases with single interaction + number of kinases with double interaction).

## Acknowledgments

We thank Corbin Jones, Brenda Temple and Anders Dohlman for their thoughtful comments on the manuscript. Funded by NIH grant R35GM118105 (to H.G.D.).

## Conflict of Interest

The authors declare no conflict of interest.

## Author Contributions

Conceptualization, methodology, validation, formal analysis, investigation, writing—original draft preparation, writing—review and editing, resources: S.L. and H.G.D.; data curation, visualization, S.L.; project administration, supervision, funding acquisition: H.G.D.. All authors have read and agreed to the published version of the manuscript.

## Footnotes

1 bioRxiv 2022.05.22.492882; doi: https://doi.org/10.1101/2022.05.22.492882

## Notes

### Competing Interest Statement

The authors have declared no competing interest.

## References

1. A. Goffeau et al., Life with 6000 genes [see comments]. Science 274, 546, 563–547 (1996).

2. K. H. Wolfe, D. C. Shields, Molecular evidence for an ancient duplication of the entire yeast genome. Nature 387, 708–713 (1997).

3. M. Kellis, B. W. Birren, E. S. Lander, Proof and evolutionary analysis of ancient genome duplication in the yeast Saccharomyces cerevisiae. Nature 428, 617–624 (2004).

4. K. P. Byrne, K. H. Wolfe, The Yeast Gene Order Browser: combining curated homology and syntenic context reveals gene fate in polyploid species. Genome Res 15, 1456–1461 (2005).

5. N. Segev, J. E. Gerst, Specialized ribosomes and specific ribosomal protein paralogs control translation of mitochondrial proteins. J Cell Biol 217, 117–126 (2018).

6. D. Solis-Escalante et al., A Minimal Set of Glycolytic Genes Reveals Strong Redundancies in Saccharomyces cerevisiae Central Metabolism. Eukaryot Cell 14, 804–816 (2015).

7. P. H. Bradley, P. A. Gibney, D. Botstein, O. G. Troyanskaya, J. D. Rabinowitz, Minor Isozymes Tailor Yeast Metabolism to Carbon Availability. mSystems 4 (2019).

8. H. F. Nijhout, F. Sadre-Marandi, J. Best, M. C. Reed, Systems Biology of Phenotypic Robustness and Plasticity. Integr Comp Biol 57, 171–184 (2017).

9. E. J. Dean, J. C. Davis, R. W. Davis, D. A. Petrov, Pervasive and persistent redundancy among duplicated genes in yeast. PLoS Genet 4, e1000113 (2008).

10. A. DeLuna et al., Exposing the fitness contribution of duplicated genes. Nat Genet 40, 676–681 (2008).

11. G. Musso et al., The extensive and condition-dependent nature of epistasis among whole-genome duplicates in yeast. Genome Res 18, 1092–1099 (2008).

12. B. VanderSluis et al., Genetic interactions reveal the evolutionary trajectories of duplicate genes. Mol Syst Biol 6, 429 (2010).

13. R. Kafri, A. Bar-Even, Y. Pilpel, Transcription control reprogramming in genetic backup circuits. Nat Genet 37, 295–299 (2005).

14. J. Stelling, U. Sauer, Z. Szallasi, F. J. Doyle, 3rd, J. Doyle, Robustness of cellular functions. Cell 118, 675–685 (2004).

15. H. Kitano, Biological robustness. Nat Rev Genet 5, 826–837 (2004).

16. M. Lynch, J. S. Conery, The evolutionary fate and consequences of duplicate genes. Science 290, 1151–1155 (2000).

17. R. Kafri, M. Levy, Y. Pilpel, The regulatory utilization of genetic redundancy through responsive backup circuits. Proc Natl Acad Sci U S A 103, 11653–11658 (2006).

18. A. Force et al., Preservation of duplicate genes by complementary, degenerative mutations. Genetics 151, 1531–1545 (1999).

19. H. Innan, F. Kondrashov, The evolution of gene duplications: classifying and distinguishing between models. Nat Rev Genet 11, 97–108 (2010).

20. A. Espinosa-Cantu, D. Ascencio, F. Barona-Gomez, A. DeLuna, Gene duplication and the evolution of moonlighting proteins. Front Genet 6, 227 (2015).

21. Y. Van de Peer, E. Mizrachi, K. Marchal, The evolutionary significance of polyploidy. Nat Rev Genet 18, 411–424 (2017).

22. J. Peter et al., Genome evolution across 1,011 Saccharomyces cerevisiae isolates. Nature 556, 339–344 (2018).

23. C. Sander, R. Schneider, Database of homology-derived protein structures and the structural meaning of sequence alignment. Proteins 9, 56–68 (1991).

24. L. A. Mirny, E. I. Shakhnovich, Universally conserved positions in protein folds: reading evolutionary signals about stability, folding kinetics and function. J Mol Biol 291, 177–196 (1999).

25. I. Mihalek, I. Res, O. Lichtarge, A family of evolution-entropy hybrid methods for ranking protein residues by importance. J Mol Biol 336, 1265–1282 (2004).

26. J. A. Capra, M. Singh, Predicting functionally important residues from sequence conservation. Bioinformatics 23, 1875–1882 (2007).

27. S. Karlin, L. Brocchieri, Evolutionary conservation of RecA genes in relation to protein structure and function. J Bacteriol 178, 1881–1894 (1996).

28. E. G. Krebs, The Albert Lasker Medical Awards. Role of the cyclic AMP-dependent protein kinase in signal transduction. JAMA 262, 1815–1818 (1989).

29. A. Varshavsky, The Ubiquitin System, Autophagy, and Regulated Protein Degradation. Annu Rev Biochem 86, 123–128 (2017).

30. L. A. Marotti, Jr., R. Newitt, Y. Wang, R. Aebersold, H. G. Dohlman, Direct identification of a G protein ubiquitination site by mass spectrometry. Biochemistry 41, 5067–5074. (2002).

31. Y. Wang, L. A. Marotti, Jr., M. J. Lee, H. G. Dohlman, Differential regulation of G protein alpha subunit trafficking by mono-and polyubiquitination. J Biol Chem 280, 284–291 (2005).

32. S. D. Cappell, R. Baker, D. Skowyra, H. G. Dohlman, Systematic analysis of essential genes reveals important regulators of G protein signaling. Mol Cell 38, 746–757 (2010).

33. M. P. Torres et al., G Protein Mono-ubiquitination by the Rsp5 Ubiquitin Ligase. J Biol Chem 284, 8940–8950 (2009).

34. G. Dixit, R. Baker, C. M. Sacks, M. P. Torres, H. G. Dohlman, Guanine nucleotide-binding protein (Galpha) endocytosis by a cascade of ubiquitin binding domain proteins is required for sustained morphogenesis and proper mating in yeast. J Biol Chem 289, 15052–15063 (2014).

35. A. Verreault, P. D. Kaufman, R. Kobayashi, B. Stillman, Nucleosomal DNA regulates the core-histone-binding subunit of the human Hat1 acetyltransferase. Curr Biol 8, 96–108 (1998).

36. A. Basu et al., Proteome-wide prediction of acetylation substrates. Proc Natl Acad Sci U S A 106, 13785–13790 (2009).

37. T. Pitti et al., N-GlyDE: a two-stage N-linked glycosylation site prediction incorporating gapped dipeptides and pattern-based encoding. Sci Rep 9, 15975 (2019).

38. S. C. Pakhrin, K. F. Aoki-Kinoshita, D. Caragea, D. B. Kc, DeepNGlyPred: A Deep Neural Network-Based Approach for Human N-Linked Glycosylation Site Prediction. Molecules 26 (2021).

39. S. B. Needleman, C. D. Wunsch, A general method applicable to the search for similarities in the amino acid sequence of two proteins. J Mol Biol 48, 443–453 (1970).

40. N. Blom, S. Gammeltoft, S. Brunak, Sequence and structure-based prediction of eukaryotic protein phosphorylation sites. J Mol Biol 294, 1351–1362 (1999).

41. J. C. Obenauer, L. C. Cantley, M. B. Yaffe, Scansite 2.0: Proteome-wide prediction of cell signaling interactions using short sequence motifs. Nucleic Acids Res 31, 3635–3641 (2003).

42. T. M. Beasley, R. E. Schumacker, Multiple Regression Approach to Analyzing Contingency Tables: Post Hoc and Planned Comparison Procedures. The Journal of Experimental Education 64, 79–93 (1995).

43. A. Agresti, An Introduction to Categorical Data Analysis (John Wiley & Sons, New York, NY, ed. 2nd, 2007).

44. G. Shan, S. Gerstenberger, Fisher’s exact approach for post hoc analysis of a chi-squared test. PLoS One 12, e0188709 (2017).

45. S. Henikoff, J. G. Henikoff, Amino acid substitution matrices from protein blocks. Proc Natl Acad Sci U S A 89, 10915–10919 (1992).

46. M. O. Collins, L. Yu, I. Campuzano, S. G. Grant, J. S. Choudhary, Phosphoproteomic analysis of the mouse brain cytosol reveals a predominance of protein phosphorylation in regions of intrinsic sequence disorder. Mol Cell Proteomics 7, 1331–1348 (2008).

47. C. R. Landry, E. D. Levy, S. W. Michnick, Weak functional constraints on phosphoproteomes. Trends Genet 25, 193–197 (2009).

48. J. Jumper et al., Highly accurate protein structure prediction with AlphaFold. Nature 596, 583–589 (2021).

49. M. Varadi et al., AlphaFold Protein Structure Database: massively expanding the structural coverage of protein-sequence space with high-accuracy models. Nucleic Acids Res 50, D439–D444 (2022).

50. D. Frishman, P. Argos, Knowledge-based protein secondary structure assignment. Proteins 23, 566–579 (1995).

51. S. Sharifpoor et al., A quantitative literature-curated gold standard for kinase-substrate pairs. Genome Biol 12, R39 (2011).

52. E. H. Fischer, D. J. Graves, E. R. Crittenden, E. G. Krebs, Structure of the site phosphorylated in the phosphorylase b to a reaction. J Biol Chem 234, 1698–1704 (1959).

53. B. E. Kemp, D. B. Bylund, T. S. Huang, E. G. Krebs, Substrate specificity of the cyclic AMP-dependent protein kinase. Proc Natl Acad Sci U S A 72, 3448–3452 (1975).

54. E. Humble et al., Non-dependence on native structure of pig liver pyruvate kinase when used as a substrate for cyclic 3’,5’-AMP-stimulated protein kinase. Biochem Biophys Res Commun 66, 614–621 (1975).

55. B. E. Kemp, E. Benjamini, E. G. Krebs, Synthetic hexapeptide substrates and inhibitors of 3’:5’-cyclic AMP-dependent protein kinase. Proc Natl Acad Sci U S A 73, 1038–1042 (1976).

56. S. B. Ficarro et al., Phosphoproteome analysis by mass spectrometry and its application to Saccharomyces cerevisiae. Nat Biotechnol 20, 301–305. (2002).

57. D. M. Phillips, The presence of acetyl groups of histones. Biochem J 87, 258–263 (1963).

58. M. LeDizet, G. Piperno, Identification of an acetylation site of Chlamydomonas alphatubulin. Proc Natl Acad Sci U S A 84, 5720–5724 (1987).

59. G. Piperno, M. LeDizet, X. J. Chang, Microtubules containing acetylated alpha-tubulin in mammalian cells in culture. J Cell Biol 104, 289–302 (1987).

60. S. C. Kim et al., Substrate and functional diversity of lysine acetylation revealed by a proteomics survey. Mol Cell 23, 607–618 (2006).

61. C. Choudhary et al., Lysine acetylation targets protein complexes and co-regulates major cellular functions. Science 325, 834–840 (2009).

62. J. Peng et al., A proteomics approach to understanding protein ubiquitination. Nat Biotechnol 21, 921–926 (2003).

63. C. Seoighe, K. H. Wolfe, Yeast genome evolution in the post-genome era. Curr Opin Microbiol 2, 548–554 (1999).

64. B. Conrad, S. E. Antonarakis, Gene duplication: a drive for phenotypic diversity and cause of human disease. Annu Rev Genomics Hum Genet 8, 17–35 (2007).

65. G. C. Conant, K. H. Wolfe, Turning a hobby into a job: how duplicated genes find new functions. Nat Rev Genet 9, 938–950 (2008).

66. B. Papp, C. Pal, L. D. Hurst, Metabolic network analysis of the causes and evolution of enzyme dispensability in yeast. Nature 429, 661–664 (2004).

67. L. Kuepfer, U. Sauer, L. M. Blank, Metabolic functions of duplicate genes in Saccharomyces cerevisiae. Genome Res 15, 1421–1430 (2005).

68. Y. Guan, M. J. Dunham, O. G. Troyanskaya, Functional analysis of gene duplications in Saccharomyces cerevisiae. Genetics 175, 933–943 (2007).

69. G. Musso, Z. Zhang, A. Emili, Retention of protein complex membership by ancient duplicated gene products in budding yeast. Trends Genet 23, 266–269 (2007).

70. J. Ihmels, S. R. Collins, M. Schuldiner, N. J. Krogan, J. S. Weissman, Backup without redundancy: genetic interactions reveal the cost of duplicate gene loss. Mol Syst Biol 3, 86 (2007).

71. X. Hu et al., Engineering and functional analysis of yeast with a monotypic 40S ribosome subunit. Proc Natl Acad Sci U S A 119 (2022).

72. R. van der Lee et al., Intrinsically disordered segments affect protein half-life in the cell and during evolution. Cell Rep 8, 1832–1844 (2014).

73. A. DeLuna, M. Springer, M. W. Kirschner, R. Kishony, Need-based up-regulation of protein levels in response to deletion of their duplicate genes. PLoS Biol 8, e1000347 (2010).

74. W. Qian, J. Zhang, Genomic evidence for adaptation by gene duplication. Genome Res 24, 1356–1362 (2014).

75. L. Hakes, J. W. Pinney, S. C. Lovell, S. G. Oliver, D. L. Robertson, All duplicates are not equal: the difference between small-scale and genome duplication. Genome Biol 8, R209 (2007).

76. R. Kafri, O. Dahan, J. Levy, Y. Pilpel, Preferential protection of protein interaction network hubs in yeast: evolved functionality of genetic redundancy. Proc Natl Acad Sci U S A 105, 1243–1248 (2008).

77. H. Yu et al., High-quality binary protein interaction map of the yeast interactome network. Science 322, 104–110 (2008).

78. H. Yu et al., Next-generation sequencing to generate interactome datasets. Nat Methods 8, 478–480 (2011).

79. M. A. Fares, O. M. Keane, C. Toft, L. Carretero-Paulet, G. W. Jones, The roles of wholegenome and small-scale duplications in the functional specialization of Saccharomyces cerevisiae genes. PLoS Genet 9, e1003176 (2013).

80. T. V. Vo et al., A Proteome-wide Fission Yeast Interactome Reveals Network Evolution Principles from Yeasts to Human. Cell 164, 310–323 (2016).

81. L. Freschi, M. Courcelles, P. Thibault, S. W. Michnick, C. R. Landry, Phosphorylation network rewiring by gene duplication. Mol Syst Biol 7, 504 (2011).

82. M. Kaganovich, M. Snyder, Phosphorylation of yeast transcription factors correlates with the evolution of novel sequence and function. J Proteome Res 11, 261–268 (2012).

83. A. van Hoof, Conserved functions of yeast genes support the duplication, degeneration and complementation model for gene duplication. Genetics 171, 1455–1461 (2005).

84. A. C. Marques, N. Vinckenbosch, D. Brawand, H. Kaessmann, Functional diversification of duplicate genes through subcellular adaptation of encoded proteins. Genome Biol 9, R54 (2008).

85. W. Qian, J. Zhang, Protein subcellular relocalization in the evolution of yeast singleton and duplicate genes. Genome Biol Evol 1, 198–204 (2009).

86. Z. Gu et al., Role of duplicate genes in genetic robustness against null mutations. Nature 421, 63–66 (2003).

87. X. Gu, Z. Zhang, W. Huang, Rapid evolution of expression and regulatory divergences after yeast gene duplication. Proc Natl Acad Sci U S A 102, 707–712 (2005).

88. G. C. Conant, K. H. Wolfe, Functional partitioning of yeast co-expression networks after genome duplication. PLoS Biol 4, e109 (2006).

89. I. Wapinski, A. Pfeffer, N. Friedman, A. Regev, Natural history and evolutionary principles of gene duplication in fungi. Nature 449, 54–61 (2007).

90. F. Monticolo, E. Palomba, M. L. Chiusano, Translation machinery reprogramming in programmed cell death in Saccharomyces cerevisiae. Cell Death Discov 7, 17 (2021).

91. T. Gera, F. Jonas, R. More, N. Barkai, Evolution of binding preferences among wholegenome duplicated transcription factors. Elife 11 (2022).

92. H. G. Dohlman, J. Thorner, M. G. Caron, R. J. Lefkowitz, Model systems for the study of seven-transmembrane-segment receptors. Annu Rev Biochem 60, 653–688 (1991).

93. S. P. H. Alexander et al., THE CONCISE GUIDE TO PHARMACOLOGY 2019/20: G protein-coupled receptors. Br J Pharmacol 176 Suppl 1, S21–S141 (2019).

94. K. F. Ostrom et al., Physiological roles of mammalian transmembrane adenylyl cyclase isoforms. Physiol Rev 102, 815–857 (2022).

95. F. Johansson, H. Toh, A comparative study of conservation and variation scores. BMC Bioinformatics 11, 388 (2010).

96. S. R. Dayhoff MO, Orcutt BC “A model of Evolutionary Change in Proteins” in Atlas of protein sequence and structure (National Biomedical Research Foundation, 1978), vol. 5, pp. 345–358.

97. H. B. Mann, D. R. Whitney, On a Test of Whether one of Two Random Variables is Stochastically Larger than the Other. The Annals of Mathematical Statistics 18, 50–60, 11 (1947).

