## Supplemental File for "Evolutionary conservation of sequence motifs at sites of protein modification"

Shuang Li and Henrik G. Dohlman

Henrik G. Dohlman


**This PDF file includes:**

Figures S1 to S8
Legends for Datasets S1 to S8

**Other supporting materials for this manuscript include the following:**

Datasets S1 to S8

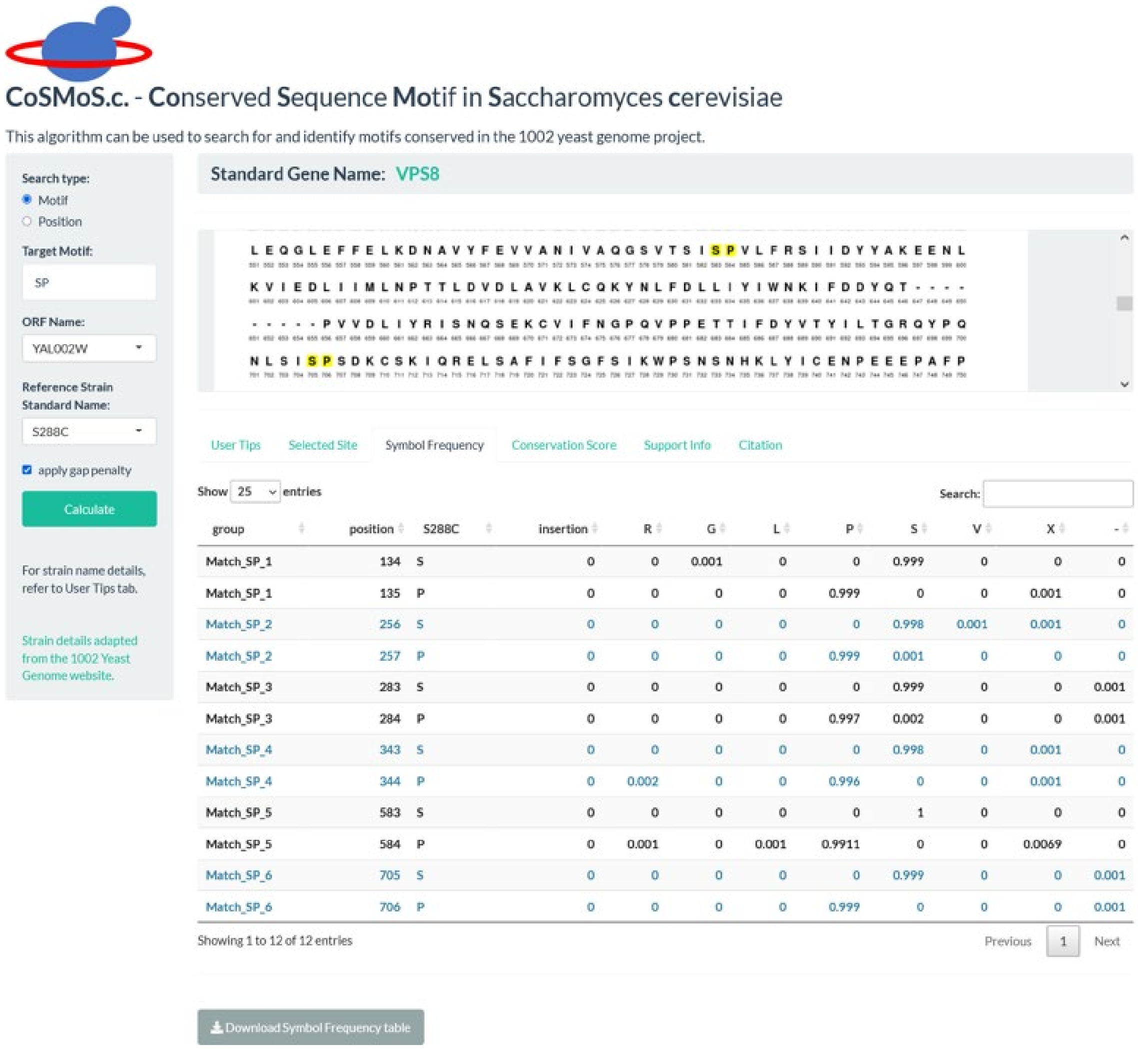


| group | position | S288C | insertion | R | G | L | P | S | V | X | - |
|---|---|---|---|---|---|---|---|---|---|---|---|
| Match_SP_1 | 134 | S | 0 | 0 | 0.001 | 0 | 0 | 0.999 | 0 | 0 | 0 |
| Match_SP_1 | 135 | P | 0 | 0 | 0 | 0 | 0.999 | 0 | 0 | 0.001 | 0 |
| Match_SP_2 | 256 | S | 0 | 0 | 0 | 0 | 0 | 0.998 | 0.001 | 0.001 | 0 |
| Match_SP_2 | 257 | P | 0 | 0 | 0 | 0 | 0.999 | 0.001 | 0 | 0 | 0 |
| Match_SP_3 | 283 | S | 0 | 0 | 0 | 0 | 0 | 0.999 | 0 | 0 | 0.001 |
| Match_SP_3 | 284 | P | 0 | 0 | 0 | 0 | 0.997 | 0.002 | 0 | 0 | 0.001 |
| Match_SP_4 | 343 | S | 0 | 0 | 0 | 0 | 0 | 0.998 | 0 | 0.001 | 0 |
| Match_SP_4 | 344 | P | 0 | 0.002 | 0 | 0 | 0.996 | 0 | 0 | 0.001 | 0 |
| Match_SP_5 | 583 | S | 0 | 0 | 0 | 0 | 0 | 1 | 0 | 0 | 0 |
| Match_SP_5 | 584 | P | 0 | 0.001 | 0 | 0.001 | 0.9911 | 0 | 0 | 0.0069 | 0 |
| Match_SP_6 | 705 | S | 0 | 0 | 0 | 0 | 0 | 0.999 | 0 | 0 | 0.001 |
| Match_SP_6 | 706 | P | 0 | 0 | 0 | 0 | 0.999 | 0 | 0 | 0 | 0.001 |

**Fig. S1.** “Symbol Frequency” shows a table of the frequency of each amino acid at all matched sites among the 1012 strains.

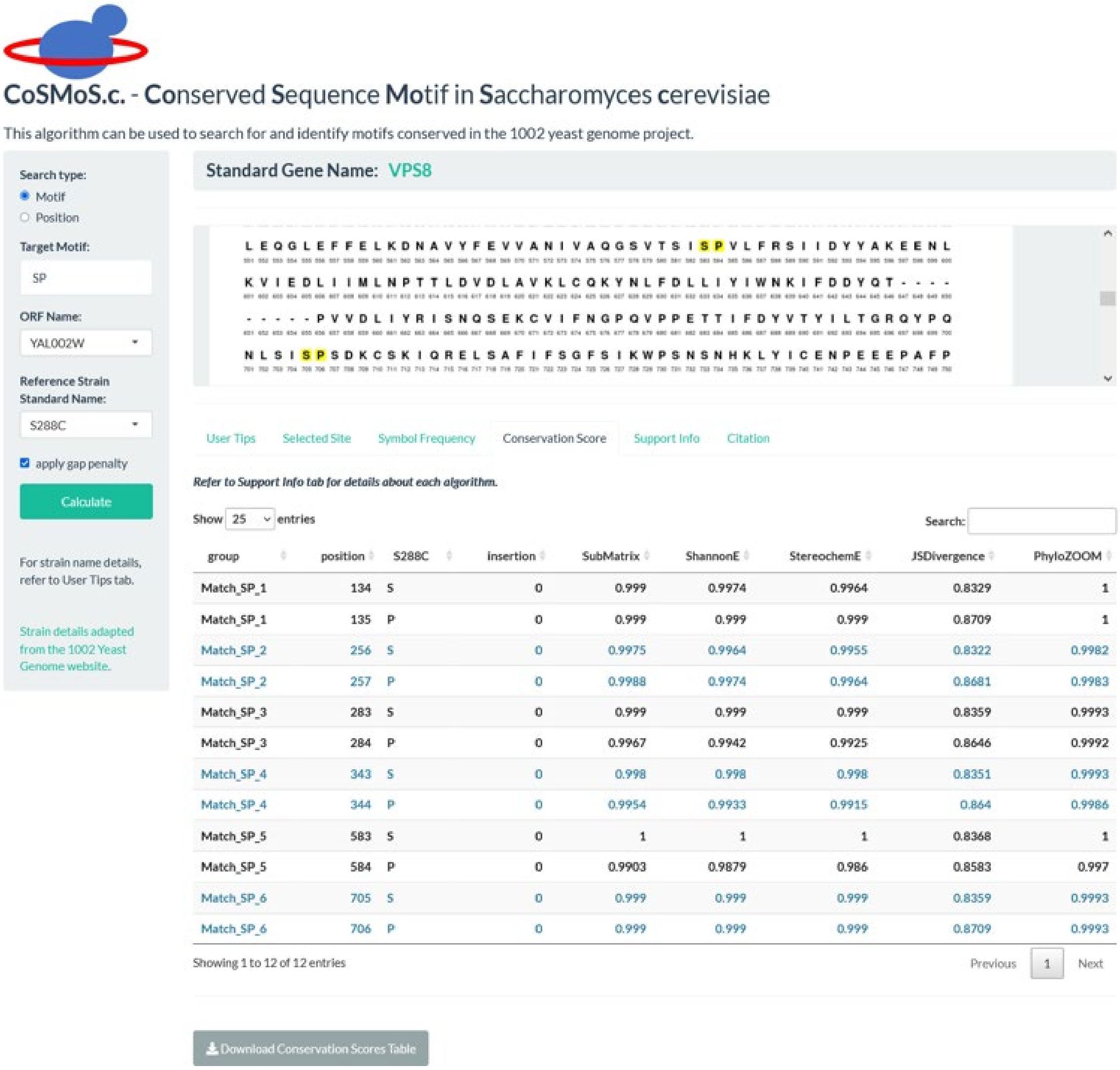


**Fig. S2.** "Conservation Score" shows a table of calculated scores for all matched sites based on five widely-used algorithms: Karlin Substitution Matrix (SubMatrix), Shannon Entropy (ShannonE), Stereochemically Sensitive Entropy (StereochemE), Jensen–Shannon Divergence (JSDivergence), and Phylogeny Entropy (PhyloZOOM).

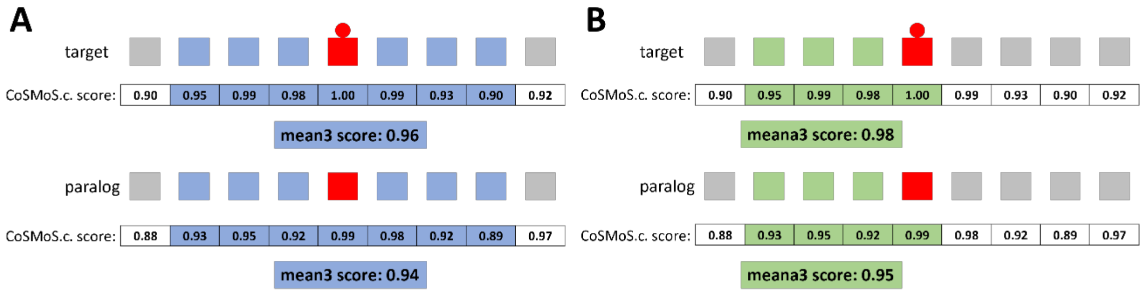


**Fig. S3.** Schematic illustrating Symmetric Average Score and One-sided Average Score. (A) Symmetric Average calculation of mean3 score for a given target protein and its paralog. (B) One-sided Average calculation of meana3 score for a given target protein and its paralog. Red circle and square shows the modified amino acid in the target protein. Red square shows the conserved amino acid in the paralog protein without modification. Amino acids and corresponding conservation scores used for meana3 score calculation are shaded in blue or green.

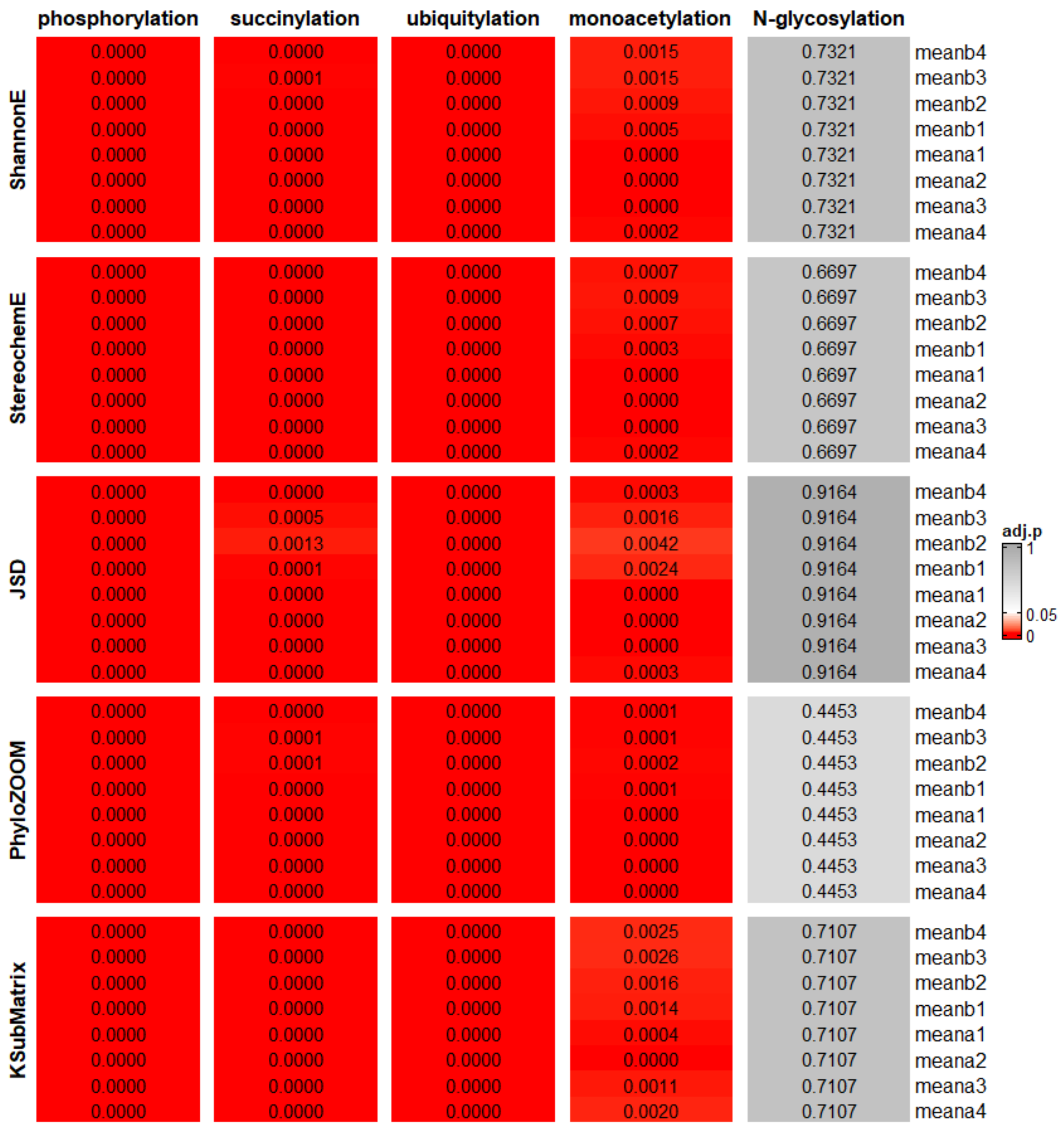


**Fig. S4.** Results of Distribution Mean Test for One-sided Average Score. Displayed as a heatmap of adjusted p values for all five algorithms with different flanking range (meanb4 to meana4, rows) for each modification type (columns). Gray, $p > 0.05$; white, $p = 0.05$; red, $p < 0.05$.

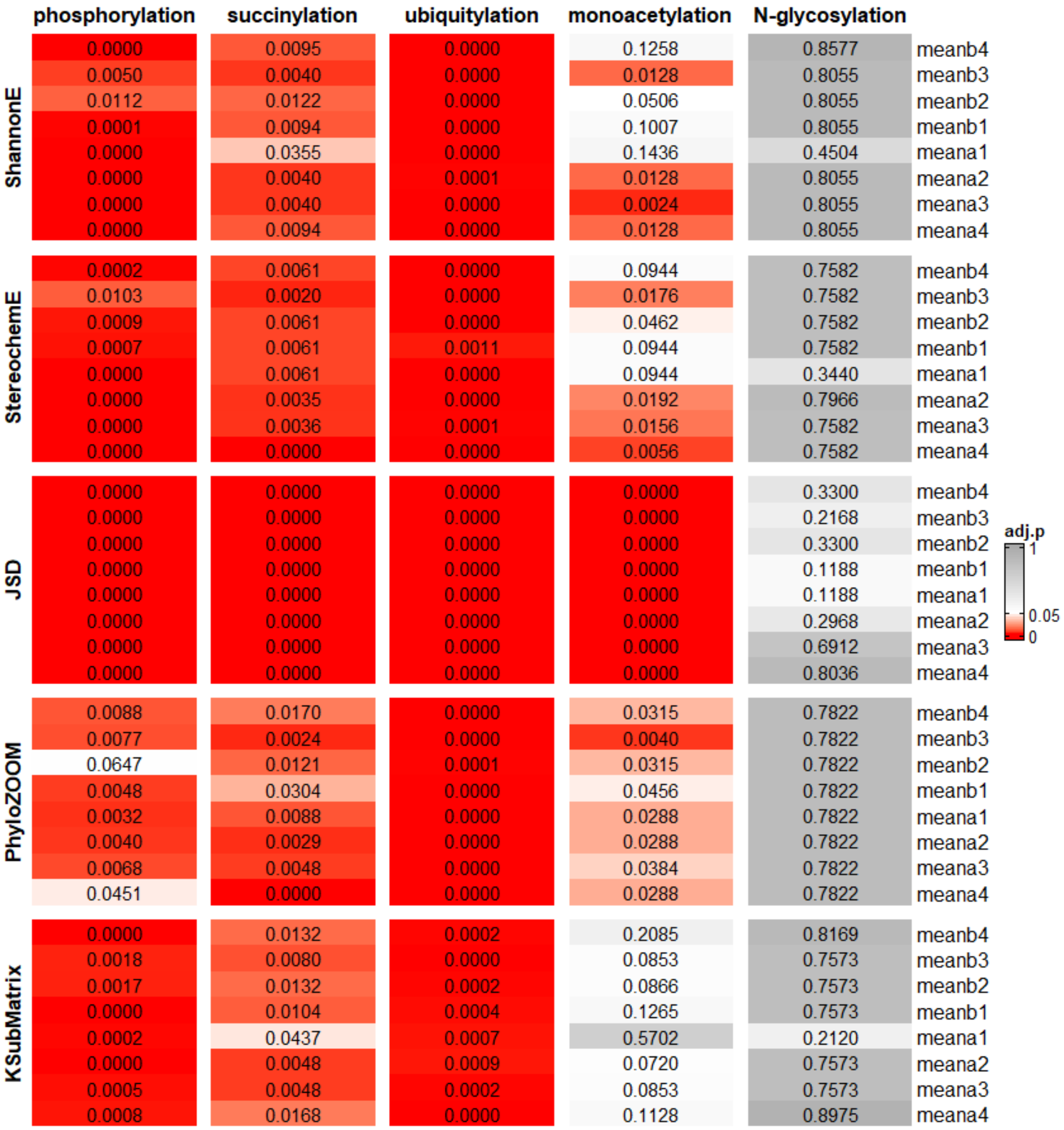


**Fig. S5.** Results of Paralog Pairing Test for One-sided Average Score. Displayed is a heatmap of adjusted p values for all five algorithms with different flanking range (meanb4 to meana4, rows) for each modification type (columns). Gray, p > 0.05; white, p = 0.05; red, p < 0.05.

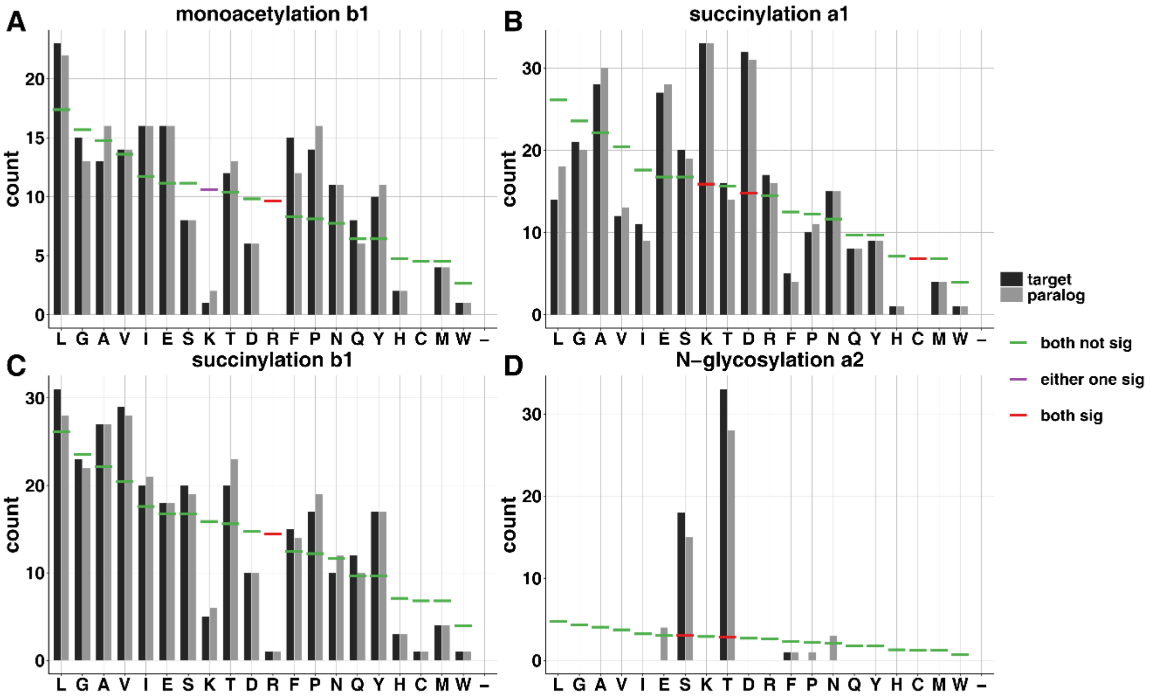


**Fig. S6.** Amino acid distribution for modified targets (black) and unmodified paralogs (grey) near sites of modifications, compared to that anticipated from BLOSUM62. A) residue before site of monoacetylation (b1); B) residue after site of succinylation (a1); C) residue before site of succinylation (b1); D) second residue after site of N-glycosylation (a2). Horizontal bars, BLOSUM62 expected amino acids frequencies. Green, neither target nor paralog is significantly different from BLOSUM62 expectation; purple, only one of target or paralog is significantly different from BLOSUM62 expectation; red, both target and paralog are significantly different from BLOSUM62 expectation.

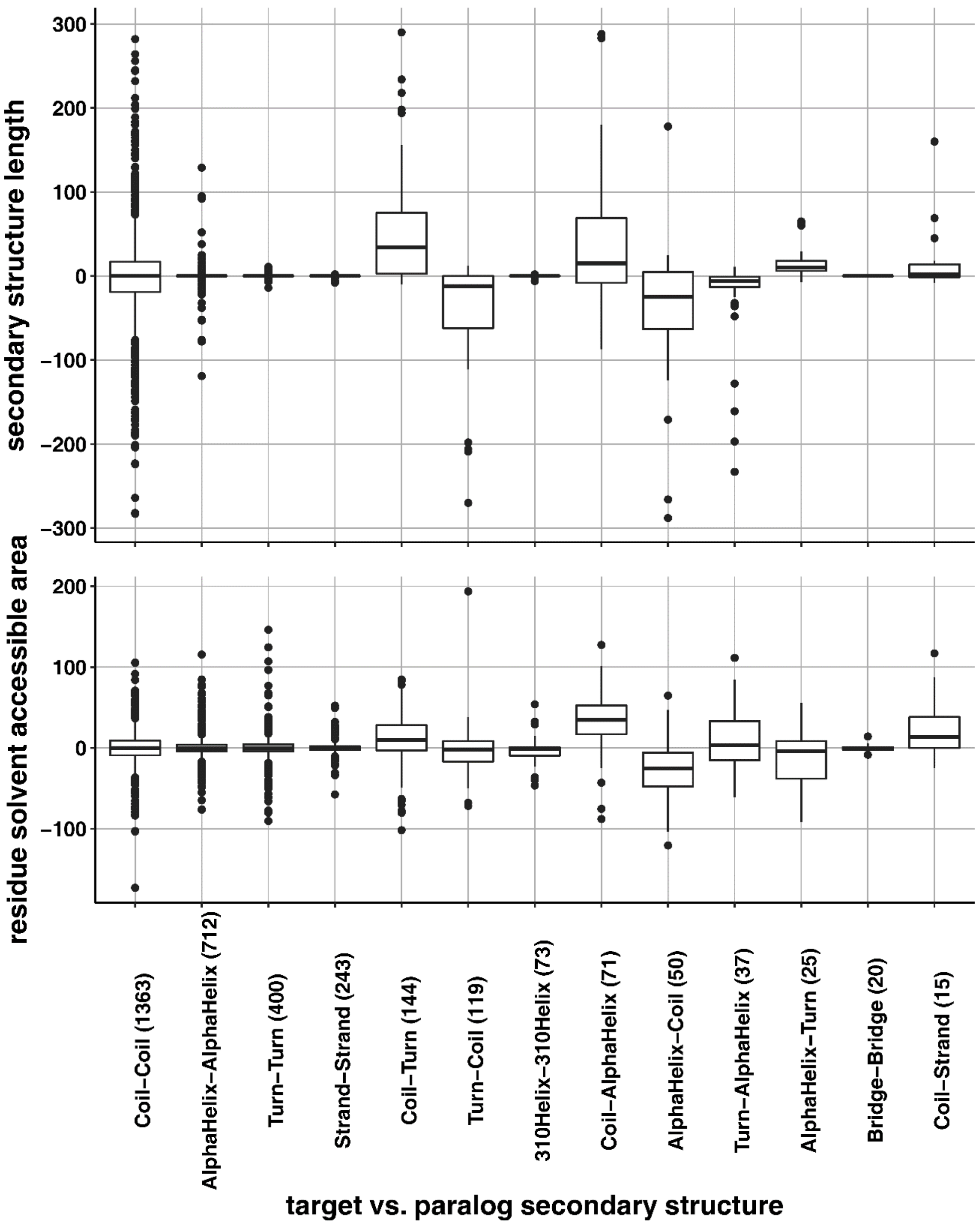


**Fig. S7.** Comparison of secondary structure properties. Box plots, differences in secondary structure length (top) and residue solvent accessible area (bottom) for each target and paralog pair. X axis, predicted secondary structure categories for each modified target site and its unmodified paralog site. Categories with fewer than 15 counts are omitted. Middle hinge, median. Lower and upper hinges, 25th and 75th percentiles respectively. The upper and lower whiskers extend from the hinge to the largest and smallest values no further than 1.5 x inter-quartile range (distance between the first and third quartiles), respectively. Data beyond the whiskers are plotted individually.

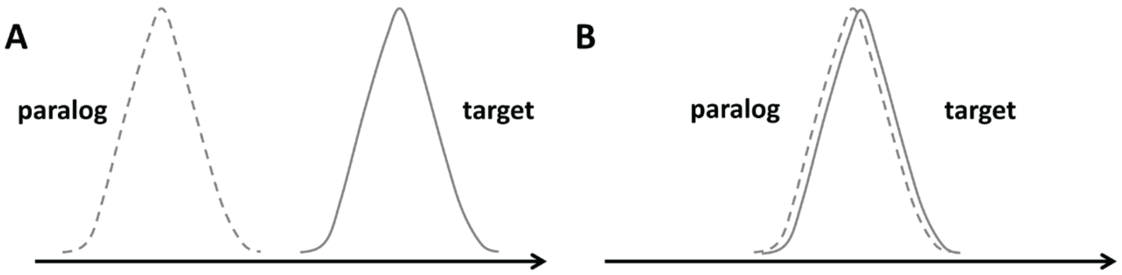


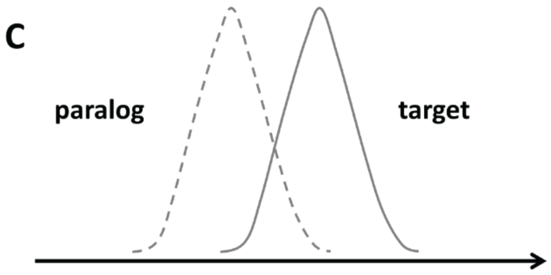


D

| | Distribution Mean Test | Paralog Pairing Test |
|---|---|---|
| A | Yes | No |
| B | No | Maybe |
| C | Maybe | Maybe |

**Fig. S8.** Schematic illustrating different target and paralog protein conservation score distributions. Shown is the conservation score distribution of target proteins (solid line) and their paralogs (dashed line). Panels A-C show different possible relationships between target and paralog conservation score distributions. Panel D shows whether Distribution Mean and Paralog Pairing tests can reveal a statistically significant difference between the two distributions.

**Dataset S1 (separate file).** Five most-common modifications among 550 paralogs. Included are ORF, gene names, modification type, modification sites, paralog ORF and gene names, Needleman aligned paralog amino acid sites and whether they are the same as the modified site (needle conserved) (Table 1).

**Dataset S2 (separate file).** Results of Distribution Mean Test for Symmetric Average Score (Fig 4).

**Dataset S3 (separate file).** Results of Paralog Pairing Test for Symmetric Average Score (Fig 5).

**Dataset S4 (separate file).** Results of Distribution Mean Test for One-sided Average Score (Fig S3).

**Dataset S5 (separate file).** Results of Paralog Pairing Test for One-sided Average Score (Fig S4).

**Dataset S6 (separate file).** Results for Chemical Similarity Average Score (Fig 6).

**Dataset S7 (separate file).** Target vs paralog integrated data including ORF name, site location, flanking sequence, modification type and secondary structure (Fig S6).

**Dataset S8 (separate file).** Paralog pairs and their interactions with kinases based on data from Yeast KID (Fig 7).

All Datasets are provided in one .xlsx file.
